# Delayed Tagging of ED-A Fibronectin-Mimetic Peptide in an RGD-Decorated Synthetic Matrix Induces Fibroblast-to-Myofibroblast Transition

**DOI:** 10.64898/2026.09.02.748971

**Authors:** Aditya S. Natu, Yinzhi Fang, Joseph M. Fox, Xinqiao Jia

## Abstract

Synthetic hydrogels with bioactive ligands have been utilized to develop 3D models to gain mechanistic insight into how discrete extracellular matrix (ECM) cues direct cell fate. While the RGD motif is ubiquitously present in healthy and diseased tissues, the <u>EDG</u>IHEL (“EDG”) sequence is present only in the extra domain A-containing fibronectin (ED-A FN), which is transiently deposited in the provisional matrix in the wound bed. Here, we explore the potential of covalently tethered EDG in conjunction with RGD to promote fibroblast-to-myofibroblast transition (FMT). Normal human lung fibroblasts (NHLFs) were maintained in bioorthogonally constructed, hyaluronan-based hydrogel (“BOHAGel”) with tethered RGD ligands. When EDG was introduced on day 0 during cell encapsulation, cellular expression of Toll-like receptor 4 (TLR4) was upregulated, and a pro-inflammatory matrix remodeling response was observed, but myofibroblast differentiation was not detected. To mimic the transition from a healthy to an injured state, we leveraged the temporal tunability of BOHAGel by supplementing cell culture media with *trans*-cyclooctene (TCO)-tagged EDG after cells were primed in the RGD environment for 8 days. As the TCO species diffused through the hydrogel, EDG was instantaneously coupled to the network through immobilized tetrazine functionalities. Delayed introduction of profibrotic EDG motifs increased mRNA levels of the myofibroblast marker (*ACTA2*), ECM proteins (*COL1A1*, *COL3A1*, *FN1*), and transforming growth factor beta1 (TGFβ1) downstream targets (*VEGFA*, *CTGF*), as well as matrix remodeling enzymes (*MMP2, TIMP1*). These changes were accompanied by the formation of αSMA stress fibers, confirming complete FMT. Delayed EDG conjugation also enhanced and reinforced α5 integrin expression. Importantly, removing the RGD signal from the gel failed to induce myofibroblast differentiation. Collectively, our results suggest that FMT depends on ligand identities and the timing of their emergence in engineered matrices.

## 1. INTRODUCTION

Many tissues in the human body are capable of regenerative self-repair following injury. During wound healing, quiescent fibroblasts become activated and undergo fibroblast-to-myofibroblast transition (FMT) through a “proto-myofibroblast” intermediate state. While proto-myofibroblasts can assemble cytoplasmic-actin stress fibers and focal adhesions, they do not express α-smooth muscle actin (αSMA).^1^ On the other hand, myofibroblasts are characterized by the neo-expression of αSMA, which is incorporated into F-actin stress fibers to generate the contractile force required for wound contraction and closure.^2^ As the primary cellular drivers of tissue repair, these cells markedly upregulate the synthesis and deposition of extracellular matrix (ECM) proteins, including fibronectin (FN) and collagen (COL). In a resolving wound, once wound closure is achieved, these cells undergo programmed apoptosis. When myofibroblast activity persists beyond wound resolution, sustained contraction and ECM deposition give rise to the stiff, scar-like tissue that characterizes fibrotic diseases.^3^ In fibrotic lungs, the branched alveolar architecture is distorted and replaced by thick, rigid, and disorganized connective tissue, leading to a severe decline in respiratory function.^4^

Because FMT is the shared initiating event in both normal wound healing and pathological fibrosis, understanding how FMT is triggered is central to controlling tissue repair outcomes. It is recognized that fibroblast activation is governed largely by signals from the surrounding ECM.^5^ Due to the inherent biocompatibility and structural similarity to the native ECM, synthetic hydrogels have been utilized to establish three-dimensional (3D) tissue models.^6^ These models have enabled molecular and cellular investigations of wound repair, particularly how matrix composition and properties alter fibroblast phenotype.^7^ To date, prevailing research on myofibroblast differentiation has focused heavily on matrix stiffness as the primary biophysical driver of fibroblast activation,^8^ neglecting the critical contributions of early biochemical signaling cascades that occur before persistent myofibroblast activation and gross tissue stiffening. There is a critical need for biomimetic 3D culture platforms that capture the evolving molecular landscape in the early stages of wound healing.

We previously reported a bioorthogonally constructed, hyaluronan (HA)-based hydrogel platform, hereafter referred to as “BOHAGel”, that permits independent, user-directed tuning of matrix properties in the presence of living cells.^9^ A cell-laden hydrogel is first prepared *via* the inverse-electron-demand Diels-Alder (IEDDA) reaction between tetrazine (Tz) and norbornene (Nb, Fig. 1A), with the former in stoichiometric excess. Several days later, a *trans*-cyclooctene (TCO)-functionalized mono- or multifunctional molecule is supplemented in the cell culture media to initiate a diffusion-controlled IEDDA reaction with tetrazine groups (Fig. 1A) dangling from the network, increasing matrix stiffness^10^ or adhesiveness^11,12^ without negatively impacting cell viability and function. Because the Tz/TCO reaction is faster than molecular diffusion through the crosslinked network, BOHAGel can be spatiotemporally modified in a physiologically relevant manner. We show that delayed introduction of the GRGDSP (RGD) motif drives epithelial-to-mesenchymal transition (EMT) in prostate cancer cells.^11^ We further demonstrate that an increase in matrix-bound RGD signal induced myofibroblast differentiation in vocal fold fibroblasts^12^ and primed prostate cancer cells for transforming growth factor beta1 (TGFβ1)-driven, SMAD2/3-mediated EMT.^13^ Crucially, hydrogels used in these studies are mechanically compliant, having an average shear elastic modulus ranging from 120 Pa up to 1,000 Pa.^10,14^

**Fig. 1:**
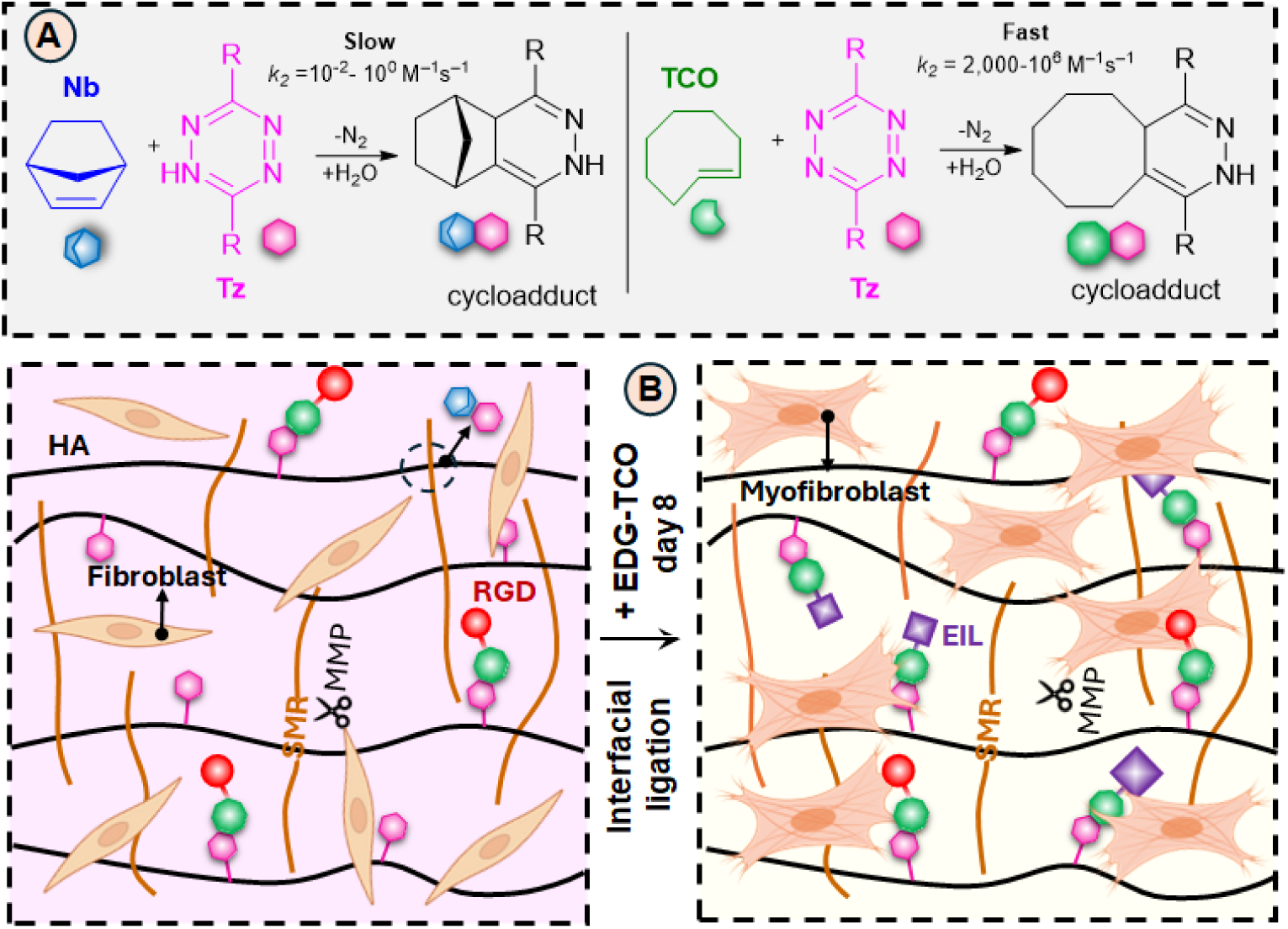
<u>B</u>io<u>o</u>rthogonal <u>h</u>y<u>a</u>luronan hydro<u>gel</u> (BOHAGel) permits sequential presentation of distinct cell-binding motifs to drive myofibroblast differentiation. (**A**) BOHAGels are fabricated using HA-Tz and a SMR-bisNb via the slow Tz/Nb cycloaddition reaction. Fibronectin-derived cell adhesive motifs, including G<u>RGD</u>SP (RGD, from domain III9/10) and <u>EDG</u>IHEL (EDG, from the ED-A domain), are covalently conjugated to the hydrogel network via the ultrafast and highly efficient Tz/TCO ligation. (**B**) The parent BOHAGel contains unreacted tetrazine groups, through which the TCO-functionalized bioactive peptide can be covalently conjugated at any time during the 3D culture. Installation of EDG signals after 7 days of cultivation in an RGD-decorated environment activates quiescent normal human lung fibroblasts to myofibroblasts.

A critical yet incompletely understood biochemical driver of fibroblast activation is the extra-domain A (ED-A)-containing isoform of fibronectin.^15^ Cellular fibronectin containing the ED-A fragment is typically absent in healthy adult tissues but is transiently deposited within the provisional matrix during active tissue repair.^16^ Full-length recombinant ED-A FN facilitates transforming growth factor beta1 (TGFβ1)-induced myofibroblast differentiation,^17^ promotes pro-fibrotic cellular response through engagement with integrins α_4β1_^18^ and α_4_β_7_,^19^ and amplifies injury-related inflammatory responses through binding with the innate immune receptor Toll-like receptor 4 (TLR4).^20^ After injury, ED-A FN appears alongside the ubiquitous RGD adhesive motifs to facilitate tissue repair.^16^ However, whether FMT depends on the temporal coordination of these signals, independently of matrix rigidity, remains to be elucidated. Although the bioactivity of ED-A FN is associated with the full ED-A domain in the context of the parent fibronectin molecule,^21^ small, well-defined sequences, such as the heptapeptide <u>EDG</u>IHEL (abbreviated as EDG) located within the C-C’ loop of the ED-A domain, have emerged as minimal motifs that recapitulate key functions of ED-A FN.^18,22–24^

Leveraging our tunable BOHAGel platform, we encapsulated normal human lung fibroblasts (NHLFs) in BOHAGels and cultivated the resulting constructs in TGFβ1-conditioned media. We presented the resident cells with RGD and EDG motifs individually, simultaneously, or sequentially. We show that RGD or EDG alone is insufficient to induce FMT. When introduced together with RGD from day 0, EDG enhances TLR4 expression and drives a pro-inflammatory fibroblast phenotype without inducing myofibroblast differentiation. Only when EDG was introduced at a delayed time point, following a period of RGD-mediated matrix engagement and TGFβ1 priming, did it drive myofibroblast differentiation (Fig. 1B). We interpret this to mean that RGD engagement and TGFβ1 stimulation first prime the fibroblasts and timely emergence of the ED-A FN-derived signal then induces the proto-myofibroblast-to-myofibroblast transition. Thus, we identify that the timing of ED-A FN emergence is a key determinant of fibroblast fate. By mimicking the transition from a healthy to an injured state, the sequential introduction of RGD and EDG signals into an otherwise soft hydrogel is sufficient to drive NHLF differentiation toward myofibroblasts without relying on elevated matrix stiffness.

## 2. METHODS

### 2.1. Synthesis of Hydrogel Precursors

Hydrogel precursors, including tetrazine-functionalized hyaluronan (HA-Tz), norbornene-tagged matrix metalloprotease (MMP)-degradable peptide VPM<u>SMR</u> (SMR-bisNb), and TCO-conjugated G<u>RGD</u>SP (RGD-TCO), were synthesized following our previously reported procedures.^9^ Using a CEM Liberty Blue peptide synthesizer (CEM, Matthews, NC), cysteine-tagged EDG peptide (EDG-SH: Ac-CEREDGIHEL-CONH_2_) was prepared on Rink amide resin (AAPPTec, Louisville, KY) employing room-temperature deprotection and coupling schemes. The peptide was cleaved with a standard trifluoracetic acid (TFA)/triisopropylsilane (TIPS)/H_2_O (95:2.5:2.5) cocktail, purified with a Teledyne ISCO reversed-phase high-performance liquid chromatography (HPLC) system (Lincoln, NE), dried on a Labconco Freezone lyophilizer (Kansas City, MO), and characterized (Fig. S1) using a Waters UPLC-MS/MS system (Xevo G2-S QTof). Peptide-bound TFA was removed by successive cycles of dissolution and drying, first in 0.1 M hydrochloric acid, then in DI water (Fig. S2). Purified EDG-SH (67 mg, 0.054 mmol) was dissolved in PBS, and the pH was adjusted to 7.4 using 0.1 M sodium hydroxide. To this solution was added 45 mg (0.081 mmol) sulfo-TCO maleimide (Broadpharm, San Diego, CA) and the mixture was stirred at ambient temperature overnight. The crude product was purified by flash column chromatography (Teledyne CombiFlash NextGen 300) using a Silica-C18 column (Biotage Sfär Bio C18, 10 g) with a gradient of 5-65% acetonitrile in water over 30 min. The product was collected by monitoring the absorbance at λ_214_/λ_280_ nm. Lyophilization yielded purified EDG-TCO (28 mg, 31% yield) as a white, fluffy solid. Product purity and molecular weight were confirmed by UPLC-MS/MS (Fig. S4). The efficiency of EDG-TCO/HA-Tz reaction was characterized using UV-vis spectroscopy (Fig. S5).

### 2.2. Fabrication of BOHAGel

Three types of BOHAGels were prepared according to procedures outlined in Fig. 2A and 2D. Stock solutions of HA-Tz (20 mg/mL) and SMR-bisNb (30 mg/mL) were prepared using 10 mM HEPES buffer containing 150 mM NaCl. RGD-TCO (15 mg/mL) and EDG-TCO (20 mg/mL) were diluted in fibroblastic basal media (FBM, Lonza, Walkersville, MD). RGD-TCO (BOHAGel-1) or RGD-TCO/EDG-TCO (BOHAGel-2) was added to the HA-Tz solution in small increments, with each addition followed by a 5-sec vortex, until the desired concentration was reached. Next, SMR-bisNb was added at a Tz/Nb molar ratio of 5/2. Forty-five minutes later, BOHAGel-1 was incubated with 4 mM EDG-TCO in PBS at 37 °C for 18 h to produce BOHAGel-3. The as-synthesized BOHAGel-2 was submerged in PBS and maintained at 37 °C overnight (∼18 h) to achieve full swelling.

**Fig. 2:**
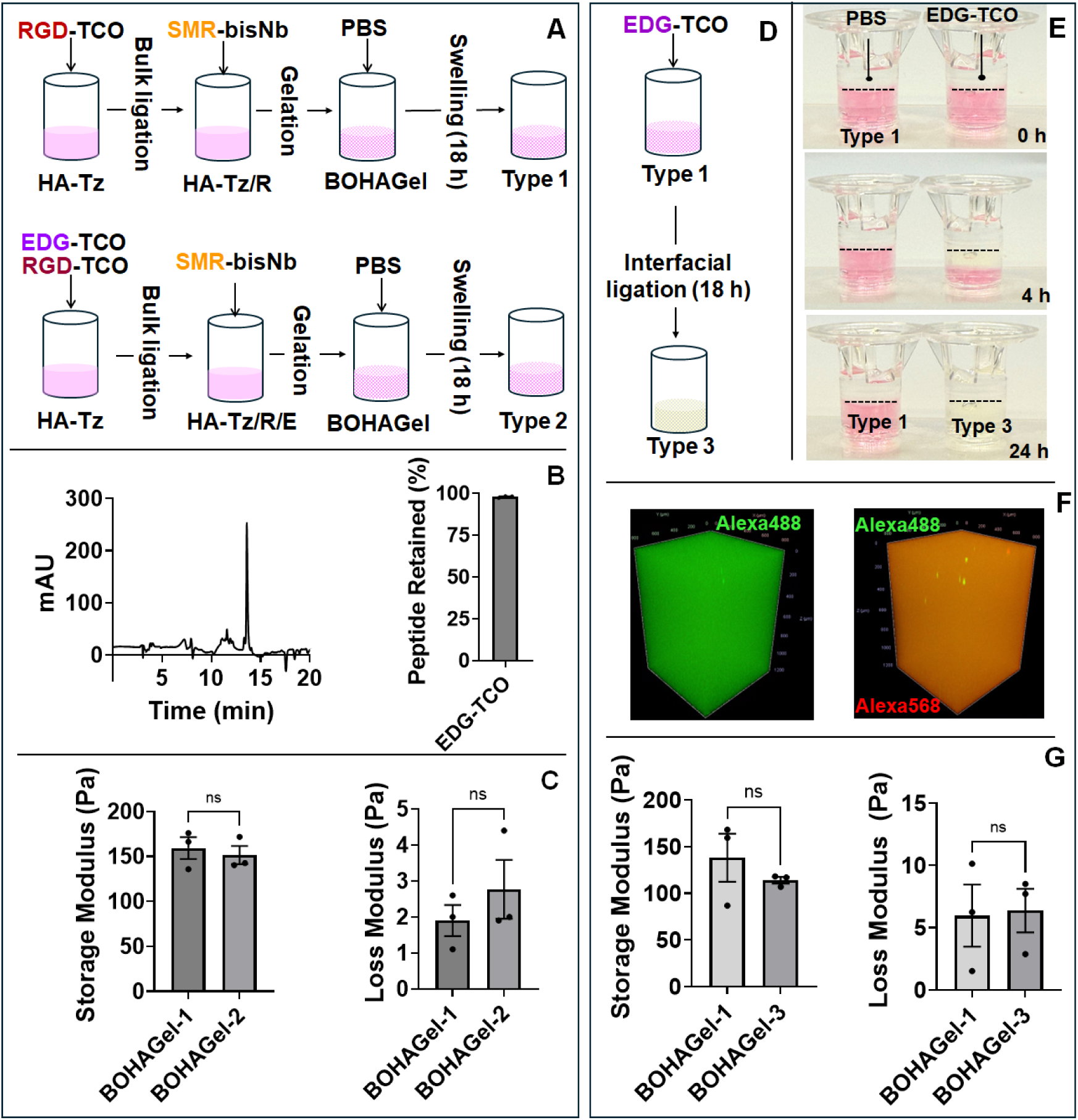
Rapid Tz/TCO reaction permits temporal introduction of bioactive peptides. (**A**) RGD and EDG were incorporated in BOHAGel via bulk conjugation. BOHAGel-1 contains 0.5 mM RGD. BOHAGel-2 contains 0.5 mM RGD and 1.0 mM EDG. (**B**) Characterization of EDG retention in BOHAGel-2. The as-synthesized gels were equilibrated in PBS for 24 h before the supernatant was aspirated for HPLC analysis (left). EDG retention (right) in the hydrogel was calculated by comparing the amount of peptide introduced and released. (**C**) Comparison of viscoelastic properties of BOHAGel-1 and 2. (**D**) Modification of BOHAGel-1 with EDG-TCO via interfacial ligation yielded BOHAGel-3 that contains 0.5 mM RGD and 2.5 mM EDG. (**E**) Time-lapse images of BOHAGel-3 (right column) 0, 4, and 24 h after incubation with EDG-TCO (4 mM in PBS). BOHAGel-1 (left column) was included for comparison. (**F**) Representative confocal images of BOHAGel before (left) and after (right) interfacial tagging. HA chains were labeled green with Alexa Fluor 488, and the EDG-TCO solution was doped with Alexa Fluor 568 (red). (**G**) Comparison of viscoelastic properties of BOHAGel-1 and 3. *n* = 3; ns: non-significant, *p* > 0.05, \*\*\*\**p* < 0.0001, determined by Student’s *t* test. Error bars represent SEM.

### 2.3. EDG Retention in BOHOGel

Type 2 BOHAGels were prepared as described above. The gel mixture (200 μL) was dispensed in a 6.5-mm cell culture insert (Cell Treat, Ayer, MA) and immersed in an additional 200 μL PBS at 37 °C for 24 h. The supernatant was then collected, and unconjugated peptide was detected at 220 nm and analyzed on an Agilent Series 1100 analytical HPLC (Agilent Technologies, Wilmington, DE) with a Phenomenex Luna 5 µm C18 100 Å column (250 mm × 4.6 mm). The standard curve (Fig. S6) was prepared by dissolving EDG-TCO in PBS at 0.05 - 1 mM. The fraction of soluble EDG peptide in the supernatant was determined following our reported method.^25^

### 2.4. Kinetics and Uniformity of Interfacial EDG Tagging

The progression of interfacial ligation was visually monitored by the disappearance of the tetrazine chromophore (pink), as previously reported.^9^ Type 1 BOHAGel was prepared, and an EDG-TCO solution (4 mM) was introduced on top. At 0, 4, and 24 h, the gel was digitally photographed. Separately, RGD-TCO containing Alexa Fluor 488-TCO (5 μM, VectorLabs, Malvern, PA) was mixed with HA-Tz before the addition of SMR-bisNb. The EDG-TCO (4 mM) solution was supplemented with Alexa Fluor 568-TCO (5 μM, VectorLabs, Malvern, PA) before being introduced to preformed BOHAGel for interfacial ligation. The modified gel was examined using a Zeiss LSM 880 microscope with a C-Apochromat 10x/0.45W objective (Carl Zeiss, White Plains, NY). Images were captured as 1.2-mm z-stacks with a 25-μm z-slice. The 3D view of the z-stack was used to generate representative images.

### 2.5. Hydrogel Viscoelasticity

The viscoelastic properties of BOHAGels were determined using a Discovery Hybrid-3 Rheometer (TA Instruments, New Castle, DE) equipped with an 8-mm parallel-plate geometry. A premade fully swollen gel disk (200 μL) was placed on the geometry, and a 0.02-N normal force was applied. Frequency sweeps were performed at 25 °C at 0.05 to 10 Hz at 5% strain. The reported storage and loss moduli are the averaged values within the linear regime from 0.05 to 0.1 Hz.^25^

### 2.6. 3D Culture in BOHAGel

#### 2.2.1. Cell maintenance

Normal Human Lung Fibroblasts (NHLFs) were purchased from Lonza Bioscience (Walkersville, MD). Cells were expanded and maintained in FBM with 5 vol% fetal bovine serum, 1 ng/mL basic fibroblastic growth factor (bFGF), 5 ng/mL insulin, and 1% (v/v) gentamicin sulfate-amphotericin. Passaging was conducted at 80-90% confluency using 0.05% (w/v) trypsin-EDTA. Trypsin was neutralized using 0.05% (w/v) trypsin soybean inhibitor (ATCC, Manassas, VA). Experiments were performed with at least three different preparations between passages 2 and 4.

#### 2.2.2. Cell encapsulation and 3D culture

NHLFs were homogeneously suspended in the hydrogel precursor mixture, prepared as described above using sterile solutions, at 3×10^6^ cells/mL. The cell suspension was transferred to 48-well glass-bottom MatTek plates (50 μL, for confocal imaging) or 8-mm cell culture inserts (200 μL, for qPCR analyses). Cellular constructs were maintained in FBM without bFGF for 24 h before the addition of TGFβ1 on day 1. Two sets of experiments (Table 1) were implemented depending on whether EDG was introduced during gelation via bulk mixing or post-gelation via interfacial ligation. In experiment A, RGD and EDG were both introduced via bulk mixing during cell encapsulation at day 0, and the resulting cultures, R_A_+E_AD0_, were maintained in FBM with 10 ng/mL TGFβ1 for a total of 14 days. Constructs prepared with RGD but without EDG served as the controls (R_A_). In experiment B, RGD was introduced via bulk mixing during cell encapsulation at day 0, and EDG was incorporated 8 days after cell encapsulation. The resulting cultures, R_B_+E_BD8_, were maintained in FBM with 5 ng/mL TGFβ1 for a total of 21 days (7 days without EDG and 14 days with EDG). Again, constructs prepared with RGD but without EDG served as the controls (R_B_).

**Table 1:** Development of NHLF-laden BOHAGel constructs.^1^. Abbreviations summarize ligand identity (R: RGD; E: EDG), day (D0 or D8) of incorporation, and culture type (A or B); ^2^. RGD was introduced via bulk mixing at 0.5 mM during cell encapsulation on day 0. ^3^. EDG was introduced via bulk mixing at 1.0 mM on day 0 or via interfacial ligation after gelation at 2.5 mM on day 8. ^4^. TGFβ1 concentration in media. ^5^. Culture duration (15 or 22 days).

| Culture ID<sup>1</sup> | RGD<sup>2</sup> |  |  | EDG<sup>3</sup> |  |  | TGFβ1<br>(ng/mL)<sup>4</sup> | Culture<br>Time (days) |
| --- | --- | --- | --- | --- | --- | --- | --- | --- |
|  | Method | Conc (mM) | Day | Method | Conc (mM) | Day |  |  |
| R<sub>A</sub> | Bulk | 0.5 | 0 | - | none | - | 10 | 15 |
| R<sub>A</sub>+E<sub>AD0</sub> | Bulk | 0.5 | 0 | Bulk | 1 | 0 | 10 | 15 |
| R<sub>B</sub> | Bulk | 0.5 | 0 | - | none | - | 5 | 22 |
| R<sub>B</sub>+E<sub>BD8</sub> | Bulk | 0.5 | 0 | Interfacial | 2.5 | 8 | 5 | 22 |

Additional control cultures generated for experiment B include R_B_+E_BD0_, where RGD was introduced via bulk mixing prior to gelation (day 0) and EDG was tagged 4 h after gelation (day 0) via interfacial ligation; E_D8_, where only EDG was introduced 8 days post-encapsulation; and NLC, where neither RGD nor EDG were introduced.

### 2.7. Cell Viability

Cellular constructs were washed with PBS before being immersed in media containing the live/dead assay reagents (ThermoFisher Scientific, City, State), which contains 4 µM calcein AM, 4 µM ethidium homodimer (EthD-1), and 40 µM Hoechst, for 30 min at 37 °C. After the media was aspirated, the constructs were thoroughly washed with PBS before being examined on a Zeiss LSM 880 confocal microscope. Three images were acquired per construct from random locations as 100 µm z-stacks with a z-axis step size of 2 µm with a 10× objective. Using ImageJ, Hoechst-positive nuclei and EthD-1-positive cells were counted from each maximum-intensity projection, and cell viability was defined as the difference between the two counts divided by the total cell count. Separately, after washing with serum-free FBM, constructs were incubated with 500 μL PrestoBlue cell viability reagent (10 vol% in FBM, Thermo Fisher Scientific) for 4 h at 37 °C. Next, 90 μL of the solution from each sample was transferred to a 96-well plate, and fluorescence was measured at 585 nm using a SpectraMax i3x Multi-Mode Microplate Reader.

### 2.8. RT-qPCR

Cellular constructs were snap-frozen in liquid nitrogen and stored at -80 °C until further processing. The frozen samples were homogenized in TRIzol (Invitrogen, Carlsbad, CA) using a pestle, then incubated in chloroform at room temperature for 5 min. After 25-min centrifugation at 16,000 rcf at 4 °C, the aqueous phase was carefully separated and purified using an RNA Clean & Concentrator-5 Kit (Zymo Research, Irvine, CA) according to the manufacturer’s protocol. RNA concentration and purity were determined using a NanoDrop 2000 spectrophotometer (NanoDrop Technologies, Wilmington, DE) based on absorbance ratios at λ_260_/λ_280_ and λ_260_/λ_230_. Following the manufacturer’s protocol, 1 μg of the extracted and purified RNA was reverse-transcribed into cDNA using the Quantitect Reverse Transcription Kit (Qiagen, Hilden, Germany). Real-time quantitative polymerase chain reaction (RT-qPCR) was performed using an Applied Biosystems 7300 real-time PCR machine (Waltham, MA). PCR reactions were prepared in a 96-well format at 20 μL by combining 2× Power SYBR green PCR master mix (Applied Biosystems), cDNA templates (0.4 ng/μL), and the target-specific primers (400 nM). All primers were synthesized by Integrated DNA Technologies (Coralville, IA). Complete primer sequences are available in Table S1. Glyceraldehyde 3-phosphate dehydrogenase (GAPDH) was used as the reference gene, and cycle threshold values were generated using 7300 System SDS RQ Study software version 1.4 (Applied Biosystems). The obtained CT values were normalized to GAPDH, and the fold changes were calculated using the ΔΔCT method. Three biological replicates are reported from three technical replicates measured in duplicate.^26^

### 2.9. Immunofluorescence

Cell-laden hydrogel constructs were washed with PBS, fixed in 4% (w/v) paraformaldehyde for 50 min at room temperature, and then washed with PBS again. Following the conditions described in Table S2, constructs were stained with the respective antibodies. Alexa Fluor 568-Phalloidin (1:400, Life Technologies, Carlsbad, CA) was included in the primary (for αSMA) or secondary antibody (for TRL4 and ITG5) solution to simultaneously label F-actin. Cell nuclei were counterstained in a DAPI solution (1:200 in 3% BSA/PBS, Life Technologies, Carlsbad, CA) for 30 min at room temperature. Fluorescent images were captured with a Zeiss LSM 880 microscope via Fast Airy scan mode with either a C-Apochromat 10×/0.45W or LD LCI Plan-Apochromat 25×0.8 Imm Korr water objective (Table S4). The brightness of each channel was uniformly adjusted in each representative image using ZEN 3.0 software.

To analyze cell morphology, the outlines of five cells per image were manually traced in ImageJ, and circularity and aspect ratio were computed for each cell using ImageJ’s built-in shape descriptors. The values for the 5 individual cells were averaged to obtain a single value per image. For each time point, a total of 27 images were analyzed (3 biological replicates, 3 hydrogel constructs per replicate, 3 images per construct). The fluorescent intensity of αSMA was quantified using a custom ImageJ macro. An F-actin mask was created to identify a F-actin defined cellular region. The integrated fluorescent intensity (IFI) of αSMA in this region was normalized to the integrated F-actin intensity in the same region to account for batch-to-batch variation in staining intensity across biological replicates.^12^ An overlap coefficient was used to quantify the extent to which the F-actin cytoskeleton was decorated by αSMA stress fibers. It was calculated as the fraction of total F-actin intensity overlapping with αSMA-positive pixels, ranging from 0 (no overlap) to 1 (complete overlap). To quantify integrin alpha 5 (ITGα5) expression, an F-actin mask was created to define the cell area. ITGα5 intensity within the area was calculated and then divided by the mask area to obtain mean fluorescence intensity (MFI).^27^ Then MFI of R_B_+E_BD8_ was normalized to R_B_. Detailed image-processing parameters, thresholding settings and mask-generation steps are provided in the supporting information.

### 2.10. Statistical Analysis

For materials characterization, quantitative results were obtained from 3 replica. Analyses of cellular constructs were performed from 3 biological repeats, each with 3 technical repeats. Error bars represent standard error of the mean (SEM) from multiple repeats. Statistical analysis was performed using Student’s t-test or one-way ANOVA. One-way ANOVA analysis was followed by Tukey’s HSD post hoc for pairwise comparison. A p-value of less than 0.05 was considered significantly different. Statistical interpretations were made using GraphPad Prism version 11.

## 3. RESULTS

### 3.1. Rapid Tz-TCO ligation allows efficient conjugation of EDG to BOHAGel

To produce TCO-functionalized EDG, cysteine was appended to the peptide C-terminus via a flexible zwitterionic ER motif, yielding EDG-SH. Adapting a standard SPPS protocol, the cleaved and purified EDG-SH was obtained as a TFA salt. The TFA salt was converted to a chloride salt by ion exchange using HCl. TFA removal was confirmed by the disappearance of the fluorine peak in the ^19^F NMR spectrum (Fig. S2), thus eliminating potential TFA-related toxicity issues (data not shown).^28^ Reacting EDG-SH with an excess of sulfo-TCO maleimide in water yielded EDG-TCO (Figs. S3, S4).

Three types of RGD-decorated BOHAGels (Fig. 2A, 2D) were prepared via tetrazine ligation at a Tz/Nb molar ratio of 5/2. To prepare type 2 gels, small aliquots of EDG-TCO were mixed with HA-Tz prior to gelation to allow for efficient coupling of ED-A FN-derived motifs to the covalent network. The TCO titration experiment (Fig. S5), implemented by monitoring the tetrazine chromophore by UV-vis spectroscopy, showed quantitative, close to 100% yield for the IEDDA reaction between EDG-TCO and HA-Tz. The resulting gels were equilibrated in PBS for 24 h at 37 °C to permit release of any free, unconjugated peptide. HPLC analysis (Fig. 2B) of the supernatant showed that >97% of peptide initially introduced was retained in the network. Because the peptide was conjugated to the network as an elastically inactive dangling chain, hydrogel mechanical properties (Fig. 2C) were not affected by the presence or absence of the EDG motifs.

Leveraging the high reactivity of TCO towards Tz, EDG was alternatively conjugated to prefabricated BAHAGel at the gel-liquid interface via a diffusion-controlled mechanism, yielding type 3 gels. Upon addition of EDG-TCO in PBS (colorless) above the hydrogel (pink due to tetrazine chromophore),^9^ the peptide progressively diffused into the matrix and reacted with residual tetrazine groups, forming a colourless Tz-TCO adduct along its path (Fig. 2E). After 24 h incubation, the tetrazine groups in BOHAGel were completely consumed, resulting in complete disappearance of the pink color. To further confirm tetrazine ligation-mediated EDG conjugation, HA-Tz was functionalized with Alexa Fluor 488, and Alexa Fluor 568-TCO was used as a fluorescent surrogate of otherwise dark EDG-TCO.^29^ Prior to interfacial ligation, the parent BOHAGel appeared homogeneously green. After interfacial ligation, BoHaGel became orange due to spatial overlap of Alexa Fluor 488 (green) and Alexa Fluor 568 (red) signals (Fig. 2F). Thus, given sufficient time, the diffusion-controlled interfacial ligation method enabled molecular-level conjugation of EDG signals throughout the hydrogel volume. Interfacial tetrazine ligation did not significantly change the mechanical properties of BOHOGels (Fig. 2G). With an average elastic modulus (G’) of 140.8 ± 28.7 Pa, BOHAGels provide a permissive and instructive environment that mimics the properties of soft connective tissues.^30^ In subsequent experiments, cell-laden EDG-decorated BOHAGel was produced either by bulk mixing on day 0 or interfacial ligation on day 8.

### 3.2. Early incorporation of EDG in RGD-containing BOHAGel does not induce FMT

The first set of experiments was implemented to assess whether simultaneous presentation of RGD and EDG motifs to NHLFs from day 0 (R_A_+E_AD0_) induces FMT; constructs with RGD signals alone (R_A_) were used for comparison (Fig. 3A). By day 15, NHLFs had adopted an elongated morphology and formed interconnected cellular networks irrespective of the presence or absence of EDG (Fig. 3B, S7). Actin was organized into bundled filaments, consistent with TGFβ1-mediated cellular adaptation in an adhesive and permissive 3D matrix.^30,31^ No qualitative differences in cell spreading or morphology were observed between the two conditions. The addition of EDG in R_A_+E_AD0_ stimulated a modest but statistically significant increase (1.09 ± 0.02-fold, *p* < 0.01) in cell proliferation, as assessed by a PrestoBlue-based assay (Fig. 3C).

**Fig. 3:**
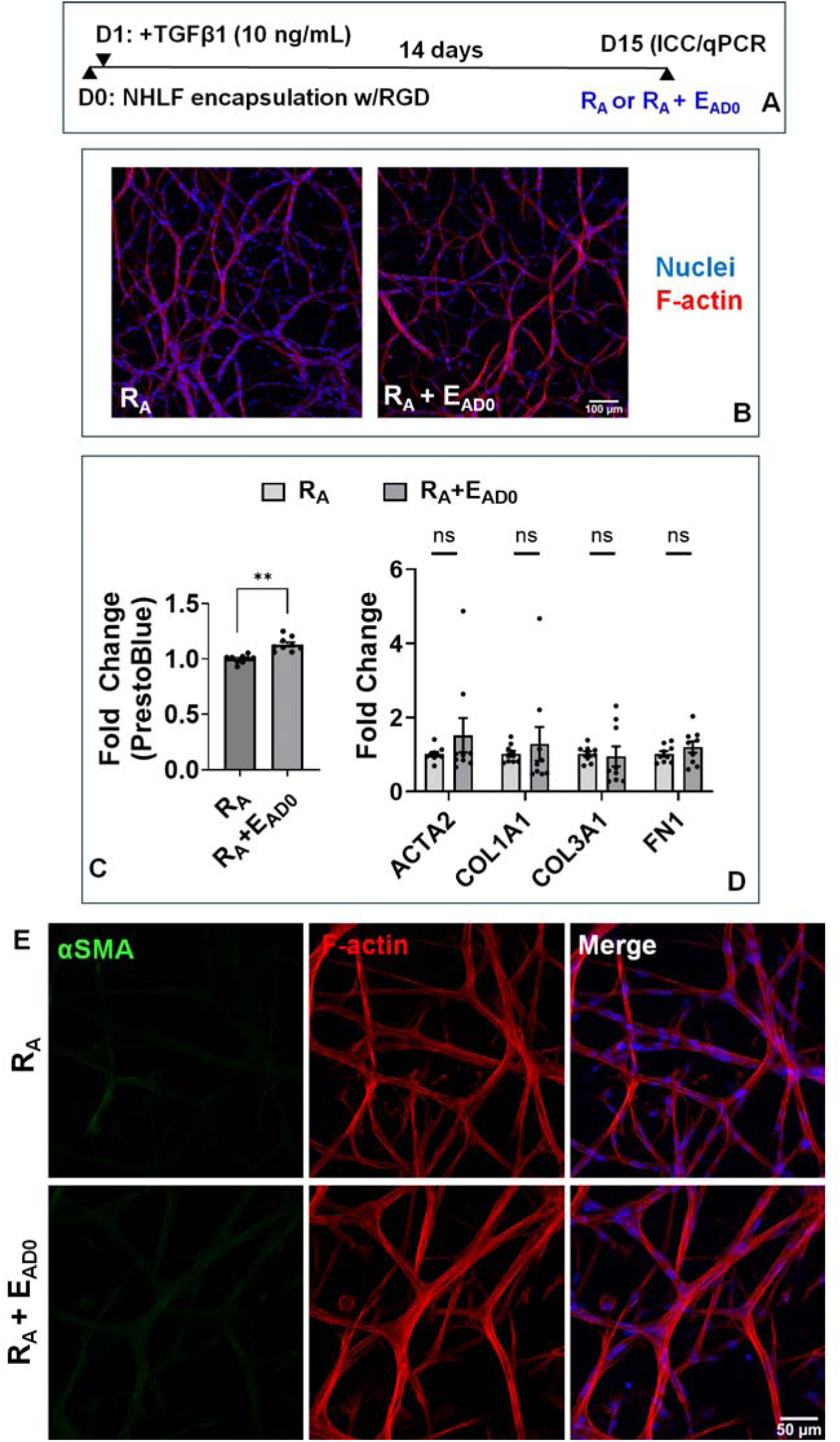
Simultaneous presentation of RGD and EDG signals from day 0 does not induce myofibroblast differentiation. (**A**) Experimental conditions and timeline. **(B)** Representative confocal images of day 15 cultures, with F-actin stained red and nuclei stained blue. Scale bar = 100 µm. (**C**) Cell viability, as measured using the PrestoBlue assay. (**D**) Expression of genes encoding myofibroblast markers. GAPDH was used as a reference gene. Expression analysis was normalized to GAPDH and reported as fold change relative to the R_A_ condition. (**E**) Representative confocal images of day 15 cultures, stained for αSMA (green), F-actin (red) and nuclei (blue). Scale bar = 50 µm. *n* = 9. Error bars represent SEM; ns: not significant, *p* > 0.05, \*\**p* < 0.01, determined by Student’s t test.

To determine whether bulk incorporation of ED-A FN epitope on day 0 at 1.0 mM is sufficient to drive myofibroblast differentiation, we next assessed the expression of canonical myofibroblast markers (Fig. 3D). Myofibroblast differentiation represents a critical phenotypic transition during wound healing, characterized by increased contractility and extracellular matrix deposition.^2^ Under the experimental conditions employed here, no significant (*p* > 0.05) changes were detected for classical myofibroblast markers including *ACTA2* (α smooth muscle actin, αSMA), *COL1A1* (collagen type 1 α1), *COL3A1* (collagen type III α1) and *FN1* (fibronectin), between the R_A_ and R_A_+E_AD0_ cultures. Immunoreactivity for αSMA in both cultures remained non-detectable (Fig. 3E)

We next examined the expression of genes encoding proteins involved in tissue remodeling and inflammatory signaling (Fig. 4A). Compared to R_A_ cultures, R_A_+E_AD0_ exhibited a significant increase in the mRNA levels of MMPs, including *MMP1* (*p* < 0.0001, 3.33 ± 0.41-fold) and *MMP9* (*p* < 0.001, 4.69 ± 1.10-fold). The expression of *MMP2*, *MMP13* and *MT1-MMP* (membrane type-1 matrix metalloproteinase), and *TIMP1* (tissue inhibitor of MMP1) remained relatively unchanged (*p* > 0.05). In addition to matrix remodelling genes, early introduction of the EDG peptide increased the transcript levels of wound healing related growth factors and inflammatory cytokines, including *HGF* (hepatocyte growth factor, *p* < 0.01, 1.96 ± 0.28-fold), *CCL2* (C-C motif chemokine ligand 2, *p* < 0.01, 1.93 ± 0.28-fold) and *CXCL10* (C-X-C motif chemokine ligand 10, *p* < 0.05, 12.98 ± 4.99-fold).

**Fig. 4:**
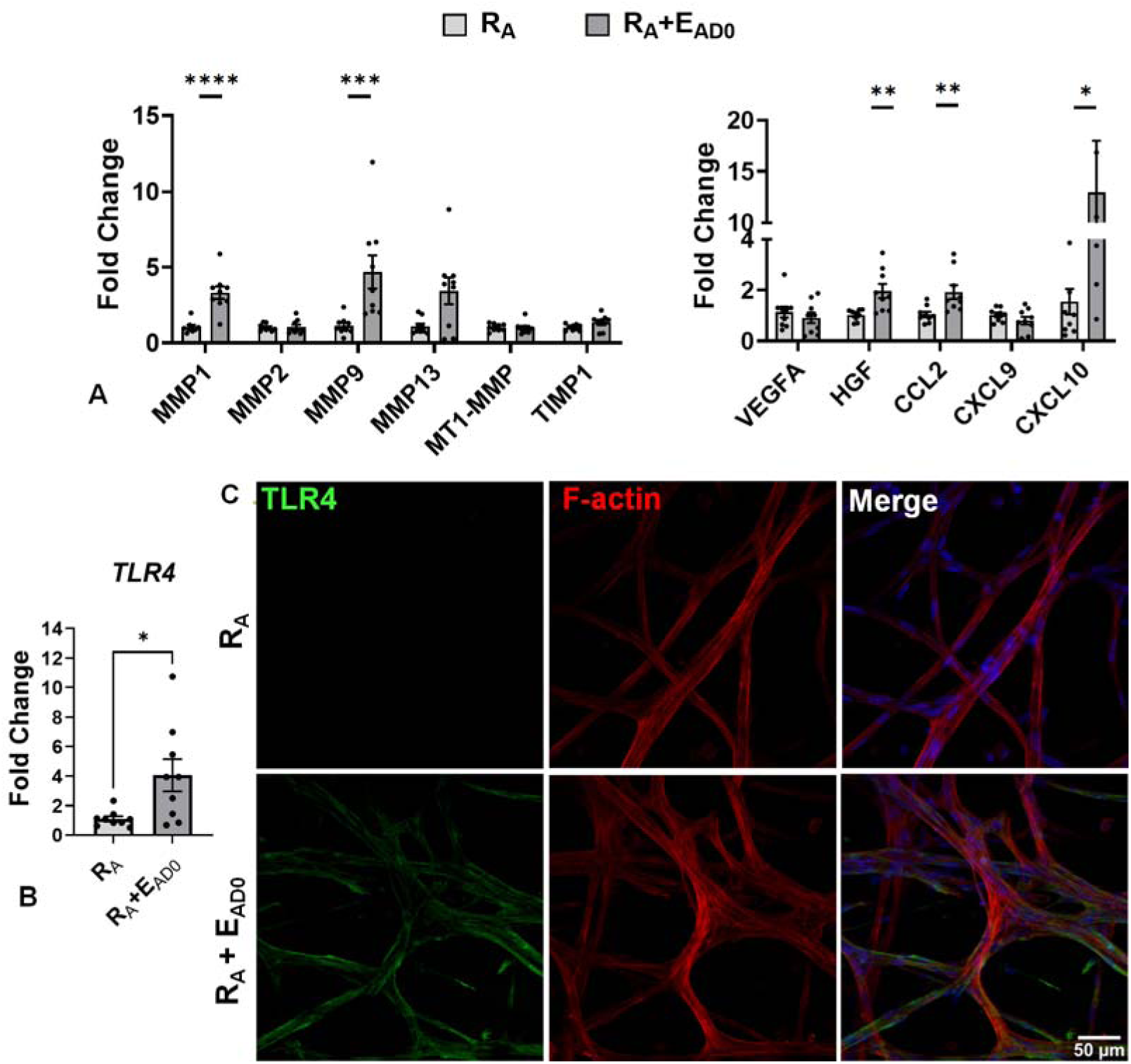
Simultaneous presentation of RGD and EDG signals from day 0 elicits a pro-inflammatory response in NHLFs. (**A**) qPCR analysis of genes encoding proteases, protease inhibitors, growth factors and cytokines. (**B**) qPCR analysis of *TLR4* expression. (**C**) Representative confocal images of NHLFs stained for TLR4 (green), F-actin (red), and nuclei (blue) in R_A_ and R_A_+E_AD0_ cultures at Day 15. Scale bars: 50 μm. All gene expression analyses were normalized to GAPDH and reported as fold change relative to the R_A_ condition. *n* = 9. Error bars represent SEM; \**p* < 0.05, \*\**p* < 0.01, \*\*\**p* < 0.001, \*\*\*\**p* < 0.0001, determined by Student’s t test.

Because ED-A FN is known to interact with TLR4 to activate innate immune signaling pathways during wound healing^32^, we queried whether the tethered EDG modulates TLR4 expression. Compared to the RGD-only controls, simultaneous presentation of RGD and EDG increased *TLR4* expression by 4.06 ± 1.09-fold (Fig. 4B) at the transcript level (*p* < 0.05). At the protein level, TLR4 signals were undetectable by immunofluorescence in cultures with RGD alone (Fig. 4C). On the other hand, R_A_+E_AD0_ cultures exhibited pronounced, robust TLR4 immunofluorescence, at the cell periphery and lamellipodia; TLR4 immunoreactivity closely mirrors the cortical F-actin network, indicating its localization on the plasma membrane where it interacts with extracellular ligands. Together, these results demonstrate that early incorporation of the EDG signal into RGD-containing BOHAGel enhances TLR4 expression and promotes a pro-inflammatory, matrix remodeling-associated transcriptional program without inducing classical myofibroblast differentiation.

### 3.3. Delayed introduction of EDG in RGD-containing BOHAGel induces FMT

To recapitulate the transition from a healthy to an early wound-repair microenvironment, we introduced the ED-A FN-mimetic peptide at a later time point after initial cell encapsulation via a rapid reaction between the diffusible EDG-TCO and the tetrazine groups dangling from the covalent network. Thus, NHLFs were encapsulated in RGD-functionalized hydrogels on day 0. From day 1 onward, constructs were maintained in media supplemented with TGFβ1 (5 ng/mL) to promote fibroblast activation. On day 8, EDG-TCO was supplemented into the media at 4 mM (Fig. 5A) and cultures were maintained under TGFβ1 stimulation for an additional 14 days to produce (R_B_+E_BD8_). Cultures with RGD only were used as controls (R_B_). The TGFβ1 concentration was reduced from 10 ng/mL to 5 ng/mL in this set of experiments to account for the longer culture period, thereby avoiding excessive TGFβ1-mediated stimulation and growth.

**Fig. 5:**
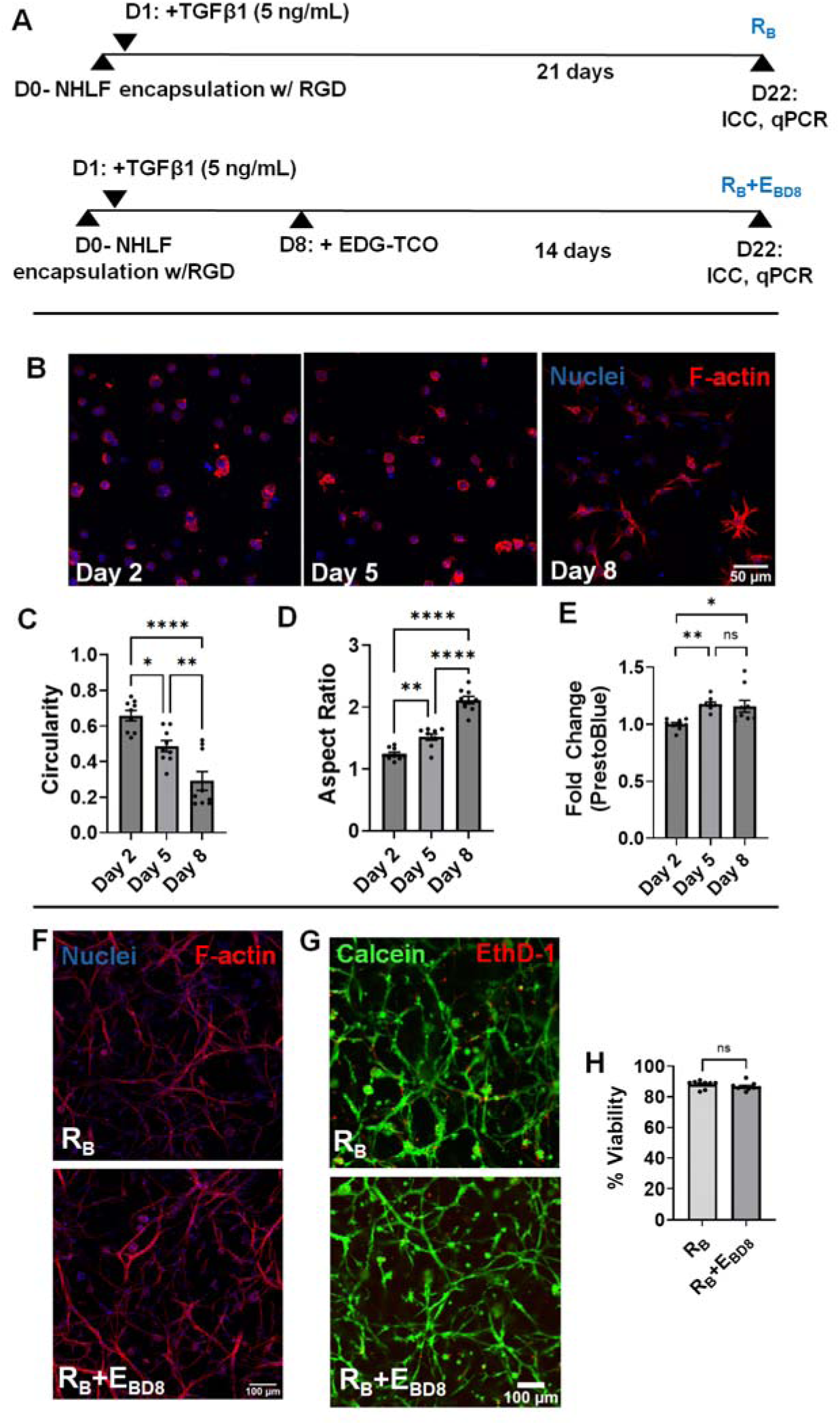
Delayed introduction of EDG in BOHAGels does not inhibit cell spreading or compromise cell viability. (**A**) Experimental conditions and timeline. (**B**) Representative confocal images of NHLFs cultured in RGD-decorated gels for 2, 5, and 8 days. F-actin and cell nuclei were stained red and blue, respectively. Scale bar = 50 μm. (**C-E**) Analysis of cell circularity (D), aspect ratio (D), and viability (E), as a function of culture time. Cell morphology (C, D) was analyzed using ImageJ software based on 27 confocal images per time point, with 5 cells per image. Cell viability (E) was assessed from via PrestoBlue assay. (**F, G**) Representative confocal images of NHLFs after F-actin/nuclei (F) and live/dead (G) staining. F-actin and nuclei are stained red and blue, respectively. Live and dead cells are stained green and red, respectively. Scale bars: 100 μm. (**H**) Quantification of cell viability based on live/dead staining. *n* = 9. Error bar: SEM; ns: not significant, *p* > 0.05; \**p* < 0.05, \*\**p* < 0.01, \*\*\*\**p* < 0.0001; determined by one-way ANOVA (C-E) or Student’s t-test (H).

Two days after cell encapsulation, NHLFs exhibited a predominantly rounded morphology with cortical actin and minimal cytoskeletal organization (Fig. 5B). By day 5, cells showed modest deviations from a spherical shape, with the emergence of short cellular extensions, although most cells remained largely rounded. By day 8, cells displayed increased spreading with the formation of small protrusions, indicating progression towards a more spread-out morphology. Quantitatively, cell circularity (Fig. 5C) progressively and significantly decreased (p < 0.0001) from day 2 (0.66 ± 0.02) to day 8 (0.29 ± 0.03). Simultaneously, the cell aspect ratio (Fig. 5D) increased significantly over time; by day 8, the average aspect ratio reached 2.11 ± 0.09. These morphological changes are consistent with cell spreading driven by integrin-mediated adhesion to RGD ligands in the matrix,^33^ in combination with TGFβ1 stimulation.^14,31^ Cell proliferation (Fig. 5E), measured indirectly using a PrestoBlue assay, showed a modest but statistically significant (*p* < 0.01) increase from day 2 to day 5; no significant change was observed from day 5 to day 8. This is consistent with enhanced cellular activity during the early stages of TGFβ1-driven cell spreading and matrix engagement, which is associated with increased metabolic demand.^34^ Thereafter, cells underwent extensive elongation overtime in both EDG-free (R_B_) and EDG-containing (R_B_+E_BD8_) cultures (Fig. 5F). Day 8 EDG ligation did not compromise NHLF viability. By day 22, over 88% of cells were viable in R_B_+E_BD8_ cultures, which was statistically indistinguishable from the R_B_ control (Fig. 5G, 5H). Together, these results demonstrate that NHLFs adapted to the 3D environment prior to EDG ligation and EDG conjugation did not hinder cells’ ability to spread in BOHAGel.

qPCR analyses revealed significant upregulations (*p* < 0.05) of canonical myofibroblast and extracellular matrix-associated genes in R_B_+E_BD8_ cultures compared to R_B_ controls (Fig. 6A). Expression of *ACTA2*, *COL1A1*, *COL3A1* and *FN1* increased by 1.54 ± 0.19 (*p* < 0.05), 1.87 ± 0.29 (*p* < 0.05), 2.11 ± 0.31 (*p* < 0.01) and 1.96 ± 0.28 folds (*p* < 0.05), respectively. The delayed EDG presentation also altered the expression of genes associated with matrix remodeling and wound-healing responses. While *MMP1* expression was significantly repressed (0.68 ± 0.03 folds, *p* < 0.05), there was a general increase in the expression of *MMP2* (1.40 ± 0.04 folds, *p* < 0.01), *VEGFA* (Vascular Endothelial Growth Factor A, 1.34 ± 0.09 folds, *p* < 0.05), *CTGF* (1.62 ± 0.12 folds, *p* < 0.05) and *TIMP1* (1.44 ± 0.15 folds, *p* < 0.05).

**Fig. 6:**
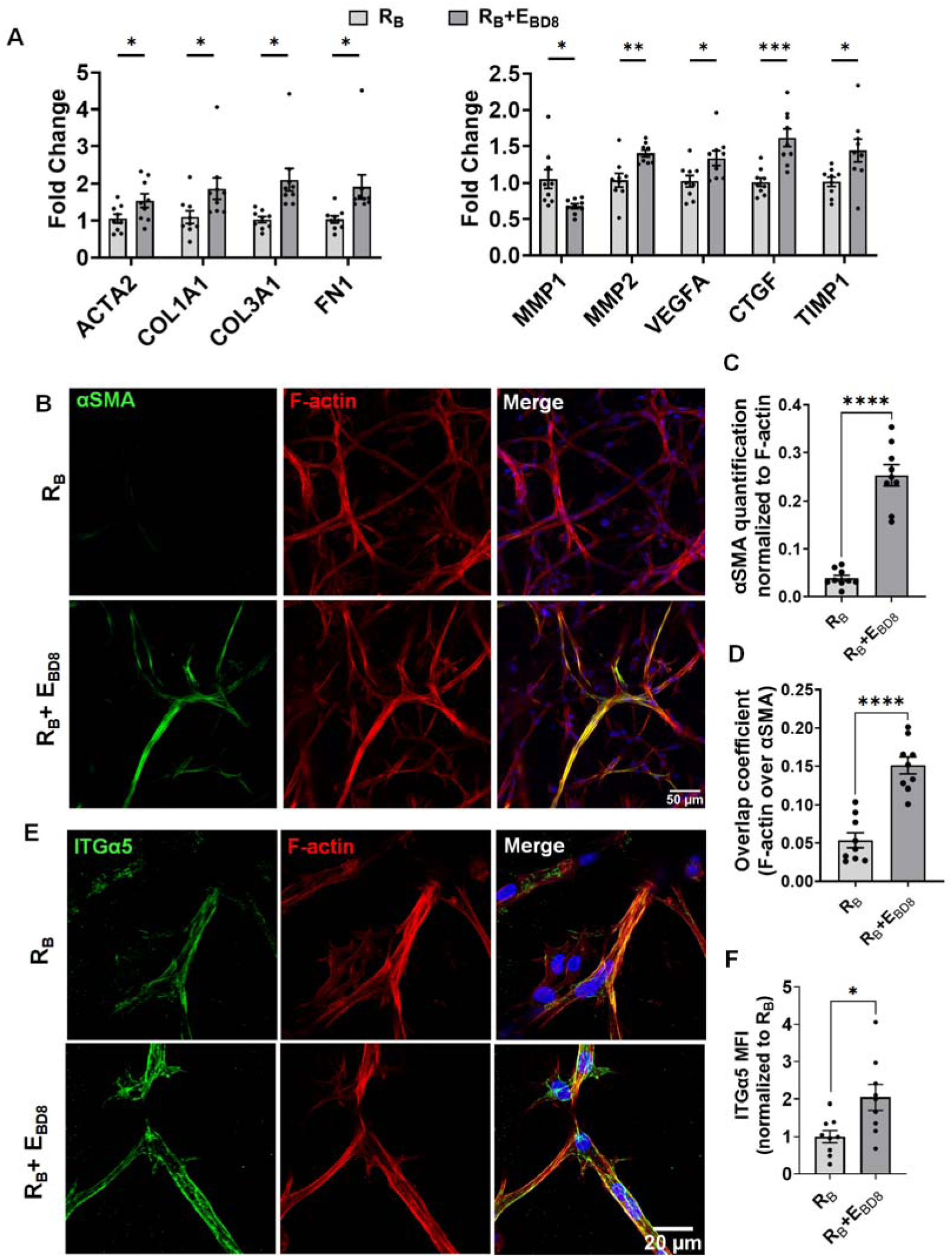
Delayed introduction of EDG in BOHAGels induces myofibroblast differentiation. (**A**) qPCR analysis of genes encoding myofibroblast markers, proteases, protease inhibitors, and growth factors. All gene expression was normalized to GAPDH and reported as fold change relative to the R_B_ condition. (**B**) Representative confocal images of 3D cultures stained for αSMA (green), F-actin (red), and nuclei (blue). Scale bar: 50 μm. (**C, D**) Quantification of αSMA immunofluorescence intensity (C) and αSMA/F-actin overlap coefficient (D, 0: no colocalization; 1: complete colocalization). (*E*) Representative confocal images of 3D cultures stained for integrin α5 (green), F-actin (red), and nuclei (blue). Scale bar: 20 μm. (**F**) Quantification of ITGα5 mean fluorescence intensity as a function of culture conditions. *n* = 9; Error bar: SEM; \**p* < 0.05, \*\**p* < 0.01, \*\*\**p* < 0.001, \*\*\*\**p* < 0.0001, determined by Student’s *t* test.

Consistent with the transcriptional changes, marked differences in αSMA immunofluorescence were observed between the two culture conditions (Fig. 6B, S8). Despite the presence of an interconnected cellular network, the R_B_ cultures showed minimal αSMA staining. In contrast, NHLFs in R_B_+E_BD8_ cultures exhibited intense αSMA staining along the cell body. Merged images showed substantial overlap between the αSMA and F-actin signals, confirming the integration of αSMA into the F-actin stress fibers. Quantitatively (Fig. 6C, 6D), this translated into a 6.3-fold increase in αSMA signal intensity (p < 0.0001) and a 2.78-fold increase (*p* < 0.0001) in αSMA/F-actin signal overlap. Together, these results demonstrate that delayed presentation of ED-A FN epitope promotes myofibroblast differentiation in NHLFs, enhancing matrix deposition, remodeling, and contractility.

To investigate whether delayed EDG introduction alters integrin expression, we examined the expression of ITGα5, the canonical RGD-binding integrin subunit, by R_B_ and R_B_+E_BD8_ cultures (Fig. 6E). While the R_B_ constructs exhibited moderate immunoreactivity for ITGa5, enhanced ITGα5 staining was observed in the R_B_+E_BD8_ cultures. In both cases, the signals were fibrillar or punctate, depending on the location. MFI quantification (Fig. 6F) confirmed a significant (2.04 ± 0.34, *p* < 0.05) increase in ITGα5 expression in R_B_+E_BD8_ samples compared to the R_B_ controls. These findings demonstrate that delayed EDG presentation enhanced cellular expression of ITGα5.

Additional control experiments (Fig 7A) were implemented to assess whether RGD-mediated matrix engagement is required and to identify the timing of EDG introduction. NHLFs were cultured in BOHAGels without RGD or EDG (NLC), without RGD but with EDG introduced on day 8 (E_BD8_), with RGD incorporated in bulk during cell encapsulation on day 0 and EDG introduced via interfacial ligation 4 hours post-gelation on day 0 R_B_+E_BD0_. In all cases, the media contained 5 ng/mL TGFβ1. By day 22, cells in NLC had relatively small cell bodies with limited spreading despite continuous TGFβ1 stimulation (Fig. 7B); cells were stained negatively for αSMA. Although a highly interconnected cellular network with prominent F-actin stress fibers was developed in E_BD8_ cultures, αSMA expression was undetectable. Interestingly, when both RGD and EDG signals were present from day 0 R_B_+E_BD0_), robust and persistent αSMA immunofluorescence was not observed even at a high EDG concentration (2.5 mM). Collectively, these results demonstrate that myofibroblast differentiation requires the basal and persistent presence of RGD, and delayed introduction of the EDG motif drives fibroblast-to-myofibroblast transition. In other words, an initial period of RGD-mediated matrix engagement and TGFβ1-driven fibroblast priming sensitizes NHLFs to subsequent EDG stimulation.

**Fig. 7:**
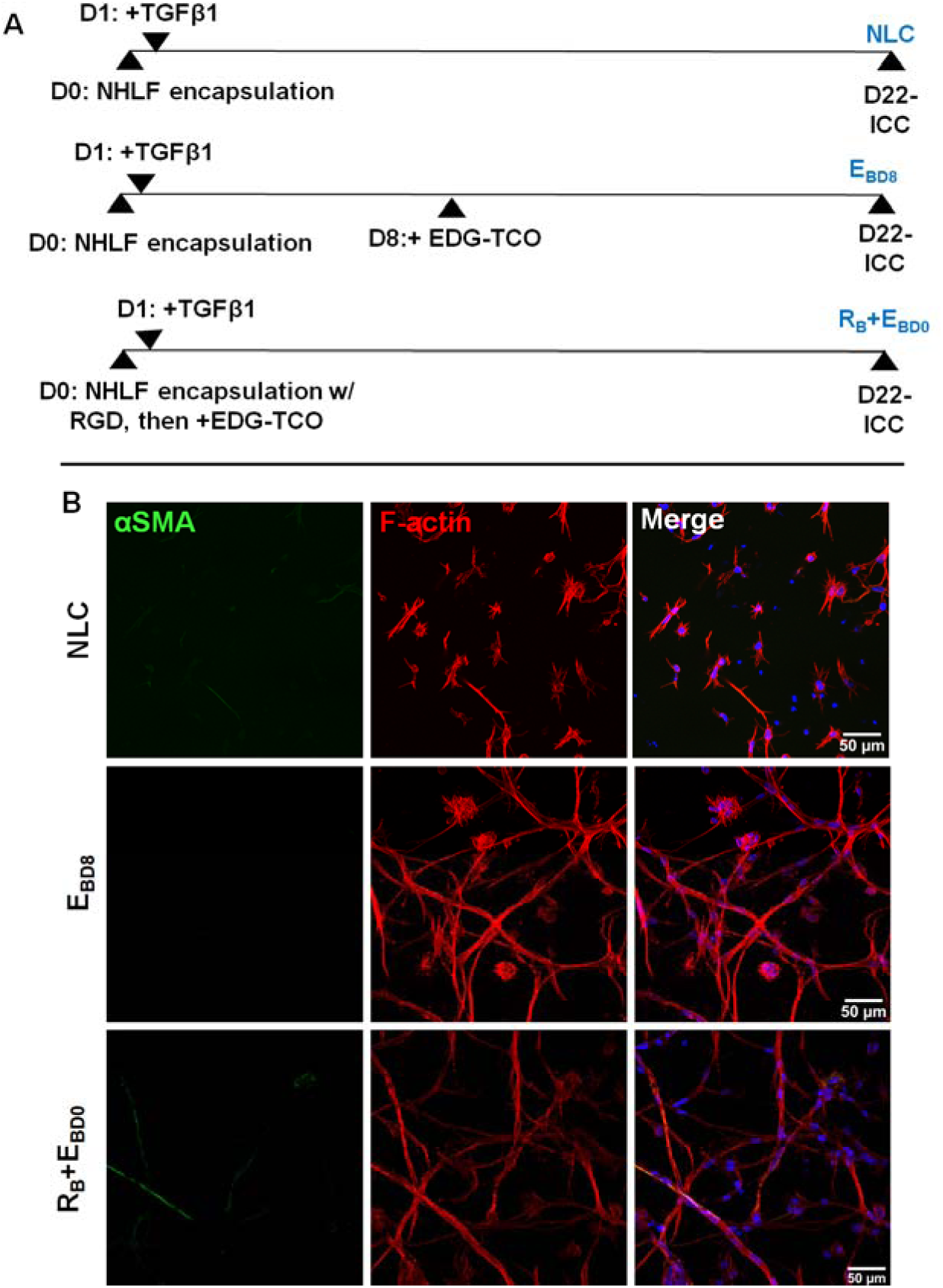
Simultaneous presentation of RGD and EDG signals throughout the entire culture period does not result in mature myofibroblast differentiation. (**A**) Experimental conditions and timeline. (**B**) Representative confocal images of NHLFs stained for αSMA (green), F-actin (red), and nuclei (blue). Scale bar = 50 µm

## 4. DISCUSSION

With an overarching goal of developing a hydrogel-based bioengineered model that closely recapitulates early stages of wound repair, we cultured NHLFs in mechanically compliant, MMP-degradable BOHAGels containing bioactive cell-binding motifs. The hydrogel platform is uniquely suitable for the targeted applications owing to its biocompatibility, bioorthogonality, and spatiotemporal tunability. It enables the simultaneous or sequential introduction of RGD and EDG motifs via the rapid, quantitative, and high-yielding IEDDA reaction between Tz and TCO.^9^ While the RGD signals are ubiquitous, the presence of ED-A FN is a hallmark of the provisional matrix of injured tissues.^35^ Through binding with integrin α_4_β_1,_^18^ α_4_β_7,_^19^ and TLR4^24^, ED-A FN synergizes with TGFβ1 to promote FMT. The EDG sequence employed here has been identified as a bioactive motif responsible for critical functions of the ED-A domain.^15,18,23,24^ Unlike the full-length protein, the short peptide epitope is straightforward to synthesize, enzymatically stable, and can be seamlessly integrated into diverse biomaterials. To our knowledge, this is the first study investigating the bioactivities of EDG in the context of 3D culture and wound healing.

For bulk incorporation, EDG-TCO was vortex-mixed with HA-Tz prior to the addition of SMR-bisNb crosslinker to ensure homogeneous distribution of the bioactive motifs in the resulting BOHAGel. Our tetrazine titration and peptide experiments confirmed that EDG-TCO reacts quantitatively with HA-Tz in solution with near-100 % efficiency. Type 2 gels with EDG concentration greater than 1 mM cannot be prepared due to the formation of insoluble precipitates. However, hydrogels with up to 2.5 mM EDG can be prepared by interfacial ligation. Our previous work demonstrates that TCO-functionalized bioactive peptides can be supplemented into cell culture media for near-quantitative reaction^11,12^ with matrix-bound tetrazines without causing significant toxicity to resident cells 4 h or 8 days post-gelation. To accelerate molecular diffusion and to ensure that the entire gel is uniformly modified, a 4 mM EDG-TCO solution was used for interfacial ligation, consuming the remaining 2.5 mM of unreacted tetrazine sites in the crosslinked network.^11,12^

We first investigated whether integrating EDG into BOHAGels with constitutive RGD can induce FMT. In agreement with previous reports,^12,14,30^ mechanically compliant, cell-adhesive, and MMP-degradable BOHAGels provide a permissive 3D environment for NHLFs to attach and extend. Under continuous TGFβ1 stimulation, NHLFs developed extensive, interconnected cellular networks throughout the matrix regardless of whether EDG is present. TGFβ1 is known to stimulate the formation of F-actin stress fibers^31^ and cell attachment is a prerequisite for TGFβ1-induced actin polymerization.^36^ The lack of significant differences in NHLF spreading in R_A_ and R_A_+E_AD0_ cultures can be attributed to the ability of RGD to engage diverse and distinct integrin heterodimers (e.g. α_5_β_1_, α_v_β_1_, and α_v_β_3_)^37^ that nucleate focal adhesions, which couple directly to the actomyosin cytoskeleton, thereby generating the traction force required for cell spreading and stress fiber assembly.^38^ In contrast, EDG exhibits strict receptor specificity, binding almost exclusively to α_4_β_1_^23^ and α_4_β_7_^19^ integrins. Importantly, integrins α_4_β_1_ regulate lamellipodial protrusion through a focal adhesion-independent mechanism.^39^ Consequently, the ubiquitous RGD-mediated mechanosignaling pathways likely dominate the matrix interface, rendering the highly restricted EDG-integrin interactions morphologically silent.

We further observed that early introduction of EDG led to upregulation of TLR4 expression at both the transcript and protein levels, consistent with the established role of the ED-A domain as an endogenous TLR4 ligand.^20,24,40,41^ This was accompanied by a coordinated pro-inflammatory and matrix-remodeling response, as evidenced by an increase in mRNA levels of proteases (*MMP1*, *MMP9*), chemokines (*CCL2* and *CXCL10*), and growth factor (*HGF*). Despite continuous TGFβ1 stimulation, early introduction of EDG to RGD-containing BOHAGel did not upregulate the expression of key myofibroblast markers, including *ACTA2*, *COL1A1*, *COL3A1*, and *FN1*. These results suggest that BOHAGel presenting RGD and EDG simultaneously from day 0 induces an inflammatory program that favors matrix turnover rather than matrix synthesis.^42^ Active turnover of collagen of this kind is a defining feature of proto-myofibroblasts, which adopt a migratory phenotype and aggressively remodel the provisional matrix instead of depositing a stable one.^43^ The pattern of MMP and chemokine induction observed here is consistent with earlier work using a recombinantly produced full-length protein containing the ED-A domain and parallels the activation of a cellular program downstream of TLR4,^32,44^ suggesting that the short EDG peptide, together with RGD, is sufficient to drive a comparable remodeling response in 3D.

Importantly, both CXCL10 and HGF have been shown to limit pulmonary fibrosis by reducing the accumulation of fibroblasts and myofibroblasts at the wound site.^45,46^ Considering the absence of αSMA-integrated stress fibers in R_A_, and R_A_+E_AD0_ cultures, as well as the R_B_+E_BD0_ control, where EDG was incorporated early (day 0) interfacially at a 2.5 mM, we speculate that the early, simultaneous presentation of the RGD and EDG motifs in BOHAGel drove NHLFs into an inflammatory, matrix-remodeling state but did not carry them through the second stage of the cascade, i.e. becoming differentiated αSMA-positive myofibroblasts. These findings are in agreement with prior reports showing that dermal fibroblasts interact with ED-A C-C’ loop-derived peptide (TYSSPEDGIHE), resulting in stress fiber formation and myosin light chain phosphorylation but without induction of αSMA.^18^

We next inquired whether sequential presentation of RGD and EDG signals in BOHAGel could stimulate robust FMT. Thus, a 7-day preculture in BOHAGel with RGD tethers was followed by a one-day incubation with EDG-TCO-conditioned media and a subsequent 14-day cultivation in the modified matrix displaying both RGD and EDG signals. TGFβ1 was present from day 1 until the culture was terminated, albeit at a lower concentration (5 ng/mL). Over the initial 7-day period, NHLFs progressively spread out, consistent with TGFβ1-stumlated, integrin-mediated cytoskeletal reorganization.^47^ By the time the EDG signal was introduced on day 8, the fibroblasts had already established stable cell-matrix interactions. Cells continued to form an extensive, interconnected cellular network in (R_B_+E_BD8_) cultures. The F-actin bundles in R_B_+E_BD8_ appeared thinner than those observed in (R_A_+E_AD0_), likely due to the lower TGFβ1 concentration in the medium.

Delayed introduction of EDG led to a marked increase in αSMA expression and its incorporation in F-actin stress fibers, unambiguously confirming the establishment of a mature, contractile myofibroblast phenotype. The upregulation of *ACTA2, COL1A1*, *COL3A1*, and *FN1* further confirms the establishment of mature myofibroblasts, capable of depositing a stable, collagen-rich provisional ECM. The concurrent decrease in *MMP1* and increase in *TIMP1* expression imply a net accumulation of newly synthesized collagen, driving matrix stabilization.^48^ Note that MMP2 cleaves of uncoiled collagen chains, rather than the fibrillar collagen.^49^ The upregulation of CTGF, widely accepted as the downstream effector of TGFβ1 signaling, suggests that EDG modulates TGFβ1 signaling.^50^ An increase in *VEGFA* expression indicates the angiogenic recruitment that accompanies wound healing.^51^

It is important to note that in the absence of either RGD or EDG (NLC), NHLFs remained relatively small, unable to assemble into networks, and exhibited minimal αSMA expression even under continuous TGFβ1 stimulation. Although cells maintained under E_BD8_ and R_D0_+E_D0_ conditions formed extended cellular networks, αSMA was not robustly integrated in the F-actin stress fibers, even at a high EDG concentration of 2.5 mM. Therefore, the timing of EDG presentation, rather than its density, determined whether NHLFs undergo FMT.

Together, our work indicates that FMT in BOHAGel requires not only the availability of RGD- and EDG-mediated cell adhesion, but also a temporal separation between them. Specifically, RGD-mediated cell-matrix engagement must be established prior to the presentation of the ED-A signal. Completion of the second stage of myofibroblast differentiation cascade requires the convergence of three inputs: elevated cytoskeletal tension, TGFβ1, and the ED-A FN-derived signal. In R_A_+E_AD0_ and R_B_+E_BD0_ constructs, both TGFβ1 and EDG were available from Day 0, but the RGD/ITGα_5_-mediated cell adhesions had not yet been materialized. Under these conditions, matrix degradation was not compensated by matrix synthesis, resulting in overall degradation of the provisional matrix, which could not support the generation of cytoskeletal tension. This tension is central to myofibroblast differentiation.^2,52^ On the other hand, without prior engagement of the RGD ligand, fibroblasts lack the cytoskeletal pre-stress required to translate the EDG signal into a differentiation response.^53^ By withholding EDG for 8 days, cells first establish RGD-dependent adhesions and enter an activated, high-tension state that can readily transition to myofibroblasts upon addition of the EDG trigger. In two-dimensional culture, ED-A containing cellular fibronectin adsorbed onto rigid substrates has been shown to induce αSMA expression in lung fibroblasts through interaction with integrin α_4_β_7_.^19^ Such substrates exceed the stiffness threshold above which αSMA is recruited to stress fibers. Thus, the cytoskeletal tension requirement is satisfied by the culture surface. In BOHAGels, the ED-A cue and cytoskeletal pre-stress could be separated due to the temporal flexibility provided by the interfacial Tz-TCO ligation system. This allowed us to resolve the FMT cascade and provide insights into how the intermediate cell state can be developed.

## 5. CONCLUSIONS

In this study, we used dynamically tunable BOHAGel to investigate the ability of a peptide mimetic of the ED-A domain of fibronectin to induce FMT from NHLFs. The 3D culture platform not only allows quantitative incorporation of multiple ECM-derived motifs but also provides control over the timing of their introduction. We demonstrate that BOHAGel containing two fibronectin-derived peptides, the ubiquitous RGD and the injury-related EDG motif, can stimulate distinct fibroblast fates that mirror successive stages of wound repair, depending on the timing of their introduction in the synthetic ECM. Simultaneous presentation of RGD and EDG from day 0 led to an inflammatory response reminiscent of the early stages of wound healing. Delayed integration of the EDG motif, after an 8-day period of RGD-mediated cell priming, stimulated robust myofibroblast differentiation, characteristic of the later matrix consolidating phase. By investigating the cooperative dynamics between the RGD and the EDG motifs, we uncovered a distinct mechanobiological checkpoint that gates myofibroblast differentiation; only fibroblasts that have first transitioned into a pre-stressed state through RGD engagement, differentiate into mature, αSMA-positive myofibroblasts. Future investigations will impart tunability not only in ligand identity but also in matrix viscoelasticity. Incorporating other injury-related cell types will enhance the physiological relevance of our model. The advanced model, combined with pharmacological inhibitors of TLR4 and integrin α5 signaling pathways, will establish the causal relationship between the timing of EDG presentation and the proto/mature myofibroblast phenotype as described here.

## ACKNOWLEDGEMENTS

This work was supported in part by the National Institutes of Health (NIH, R01DE029655, R01DC014461) and the National Science Foundation (NSF, DMR 2243648). The authors acknowledge the use of facilities and instrumentation supported by NSF through the University of Delaware Materials Research Science and Engineering Center (DMR 2011824). Microscopy access was supported by grants from the NIH-NIGMS (P20 GM103446), the NSF (IIA-1301765), and the State of Delaware. This research also benefited from the BioStore data management resource at the University of Delaware Bioinformatics Data Science Core (RRID: SCR_017696), supported by an NIH shared instrumentation grant (NIH S10 OD028725) and the Delaware INBRE (P20 GM103446). We acknowledge the Bioimaging Center at Delaware Biotechnology Institute for support on confocal imaging and image analysis. We also thank Genzyme-Sanofi for generously providing HA.

## DATA AVAILABILITY STATEMENT

The data that support the findings of this study are available from the corresponding author upon reasonable request.

## CONSENT FOR PUBLICATION

All participants provided written informed consent to the study.

