## Supporting Information for "Delayed Tagging of ED-A Fibronectin-Mimetic Peptide in an RGD-Decorated Synthetic Matrix Induces Fibroblast-to-Myofibroblast Transition"

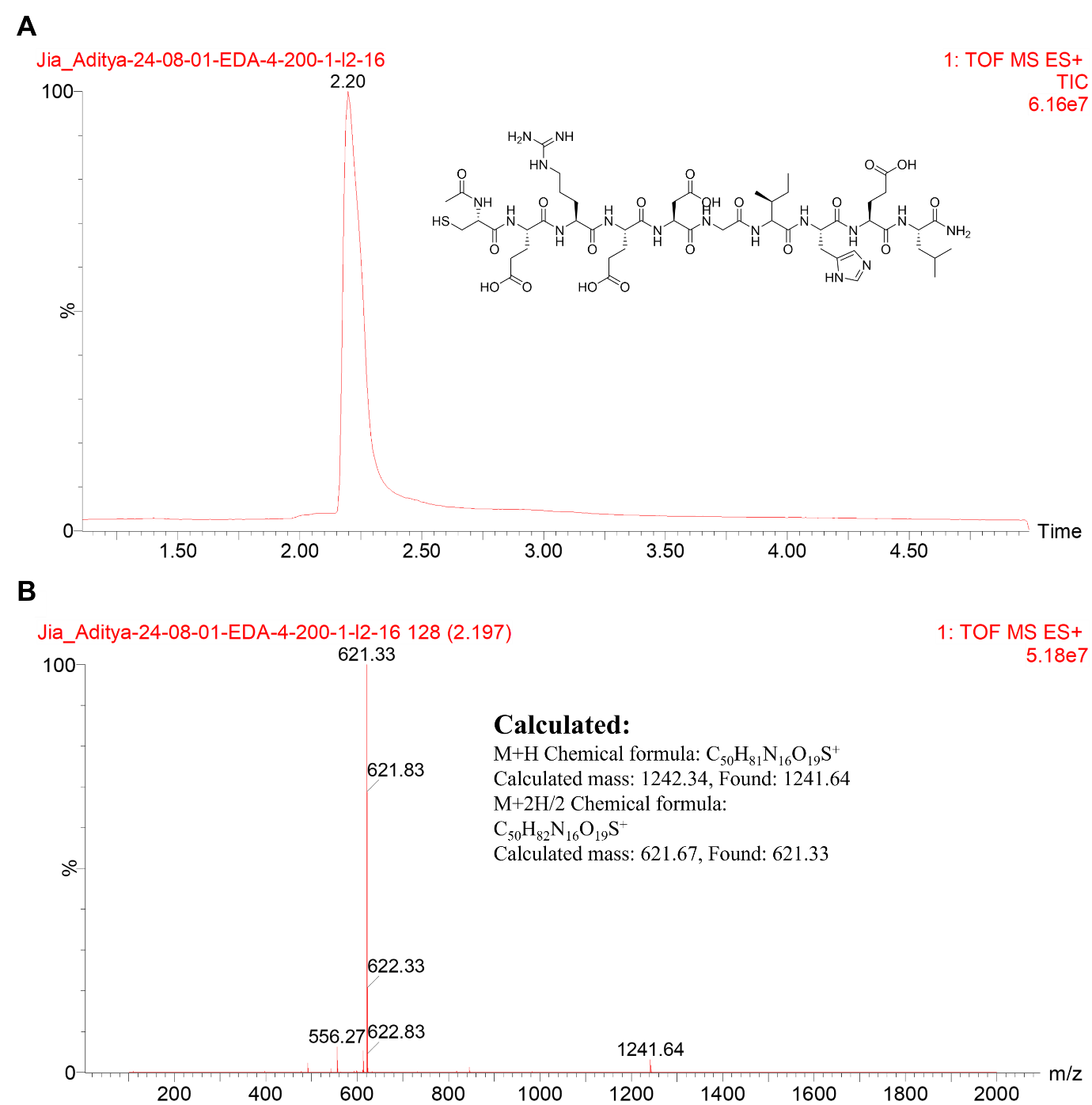

**Fig. S1:** UPLC trace (A) and mass spectrum (B) of purified EDG-SH.

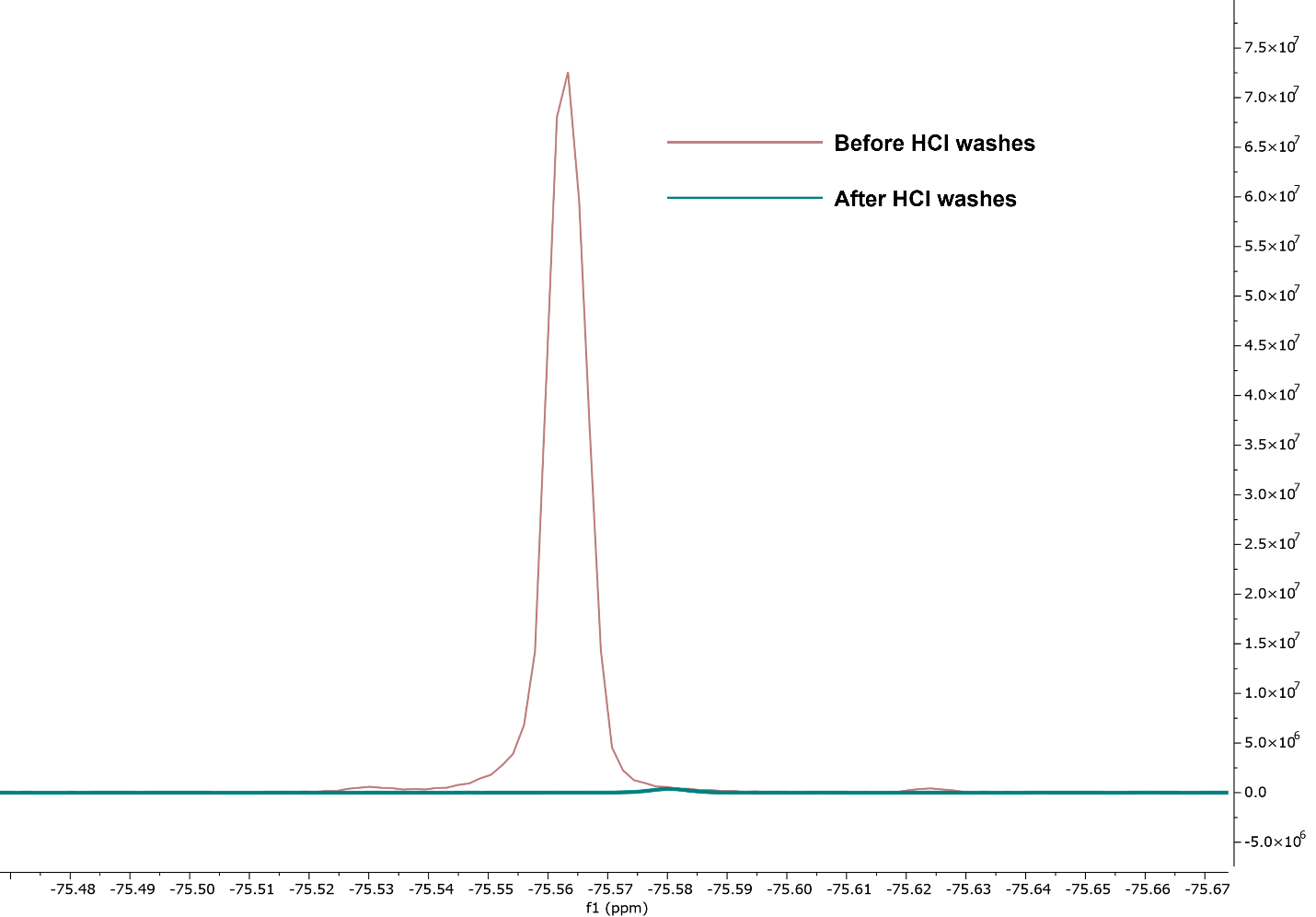

**Fig. S2:** ^19^F NMR (D_2_O with 10 vol% H_2_O, 400 MHz) of the EDG-SH peptide (20 mg/mL) before and after washing with HCl. The disappearance of the singlet peak at δ = -75.5 ppm confirmed the removal of TFA.

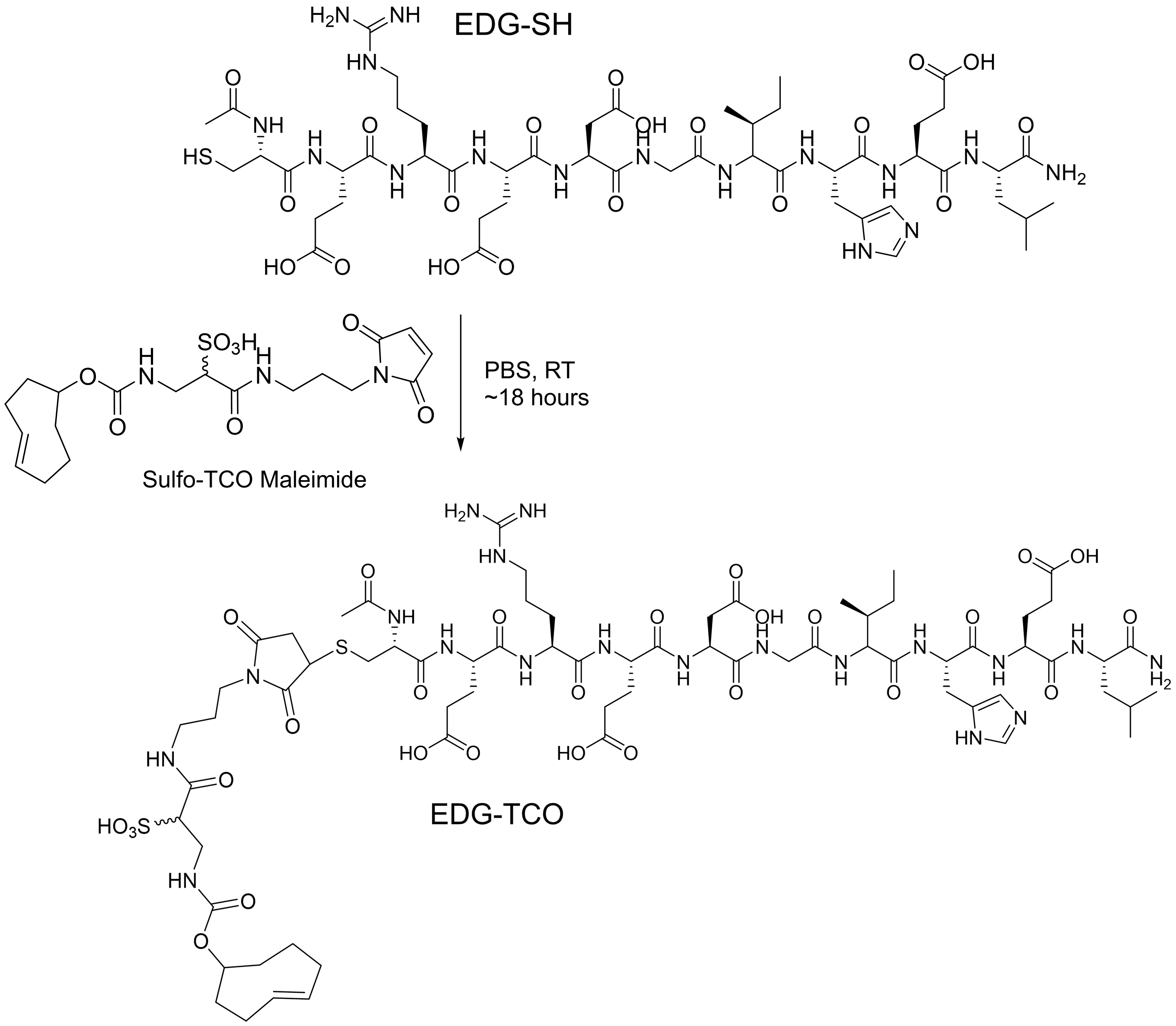

**Fig. S3:** Synthesis of EDG-TCO using sulfo-TCO maleimide and EDG-SH.

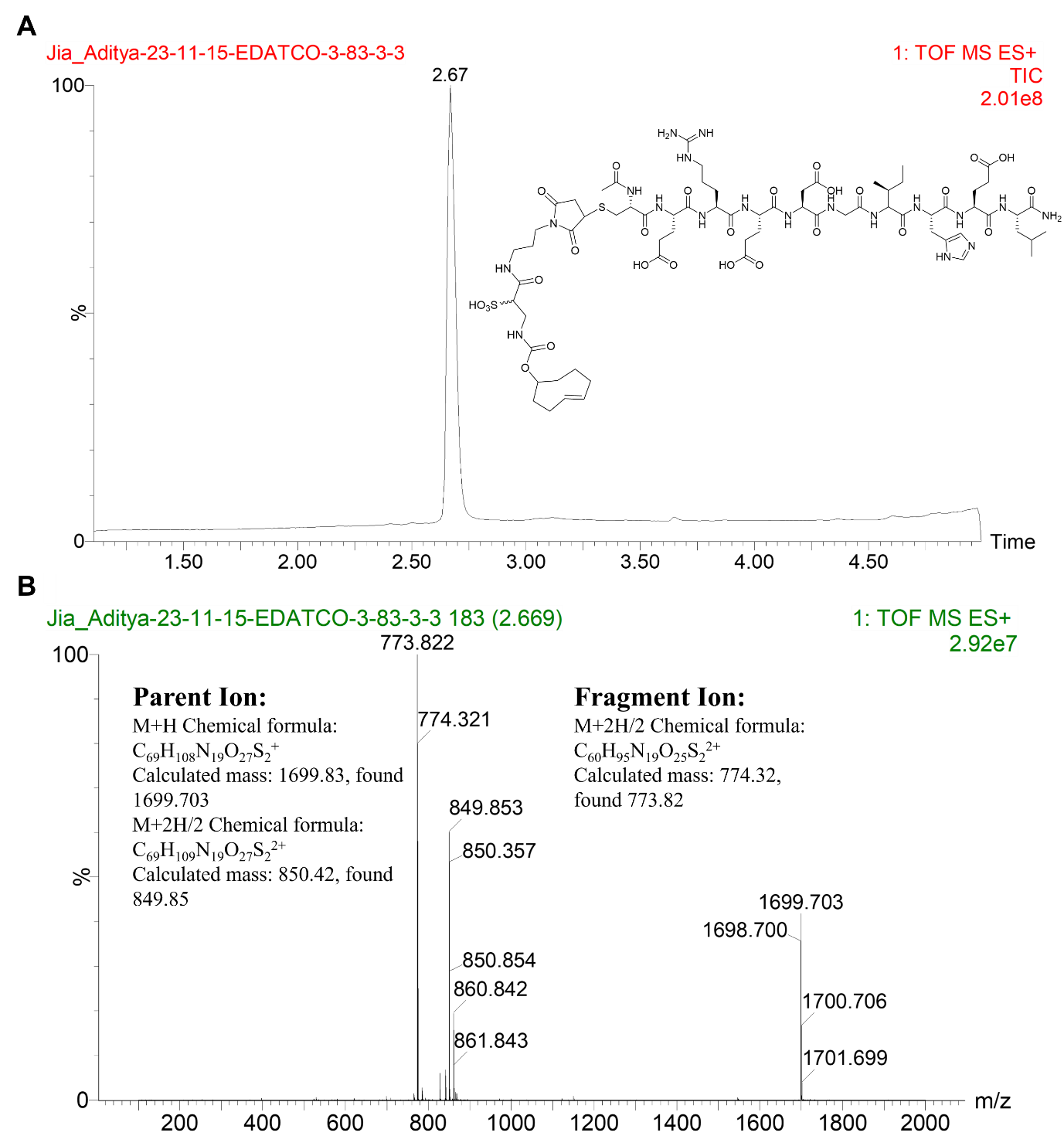

**Fig. S4:** UPLC trace (A) and mass spectrum (B) of purified EDG-TCO.

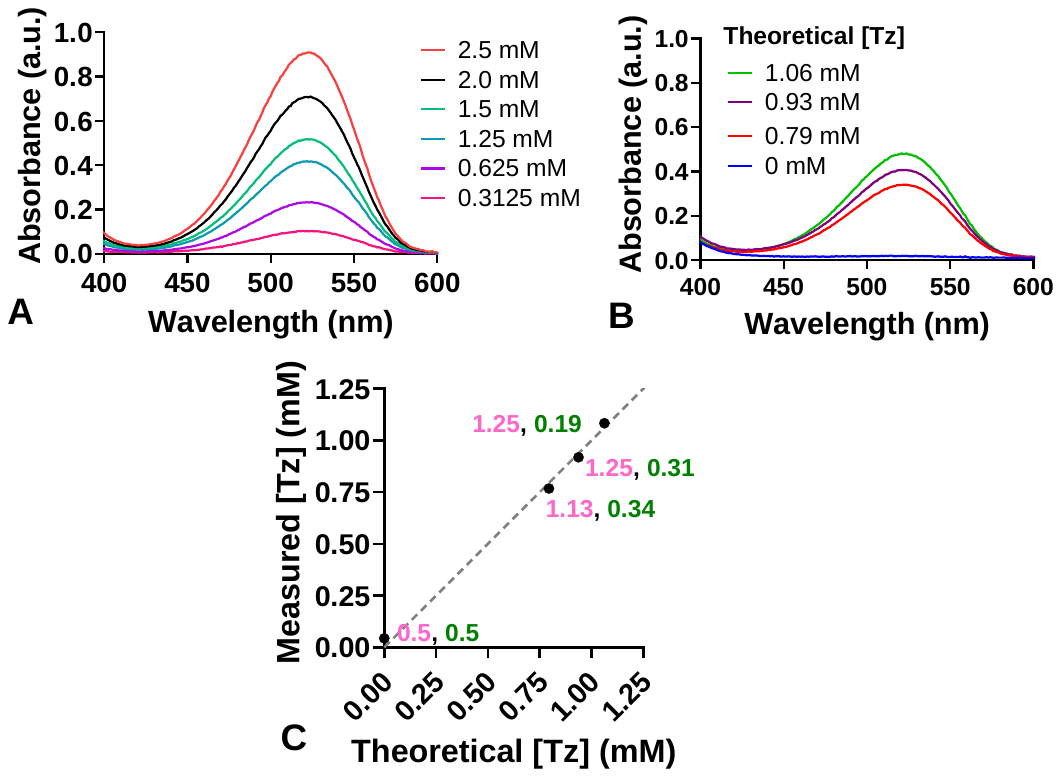

**Fig. S5:** Determination of the efficiency of the reaction between HA-Tz and EDG-TCO in PBS at pH 7.4 by UV-vis spectroscopy. (**A**) UV-vis absorbance at 534 nm, based on the methyl phenyl tetrazine chromophore, as a function of Tz concentration. Standard curves were constructed by monitoring Tz absorbance using HA-Tz (22 mol% Tz) solutions at varying Tz concentrations. The molar absorptivity coefficient (411.98 Lmol^-1^s^-1^) was calculated using Beer-Lambert’s law and the previously reported molar absorptivity coefficient for Tz at 267 nm.^1^ (**B**) UV-vis absorbance of various reaction mixture. Insert indicates theoretical Tz concentration in the mixture after the IEDDA reaction. HA-Tz and EDG-TCO were dissolved in PBS at 5 mM and 2.5 mM, respectively. The solutions were combined at different Tz/TCO molar ratios. Two minutes later, solution absorbance at 534 nm was measured using UV-vis spectroscopy. The tetrazine concentration after the reaction was calculated using the molar absorptivity at 534 nm. The theoretical values were determined by subtracting the [TCO] from [Tz] in feed. (**C**) Comparison of theoretical and experimental Tz concentrations after the IEDDA reaction. Numbers along the trend line indicate Tz (pink) and TCO (green) concentrations in feed.

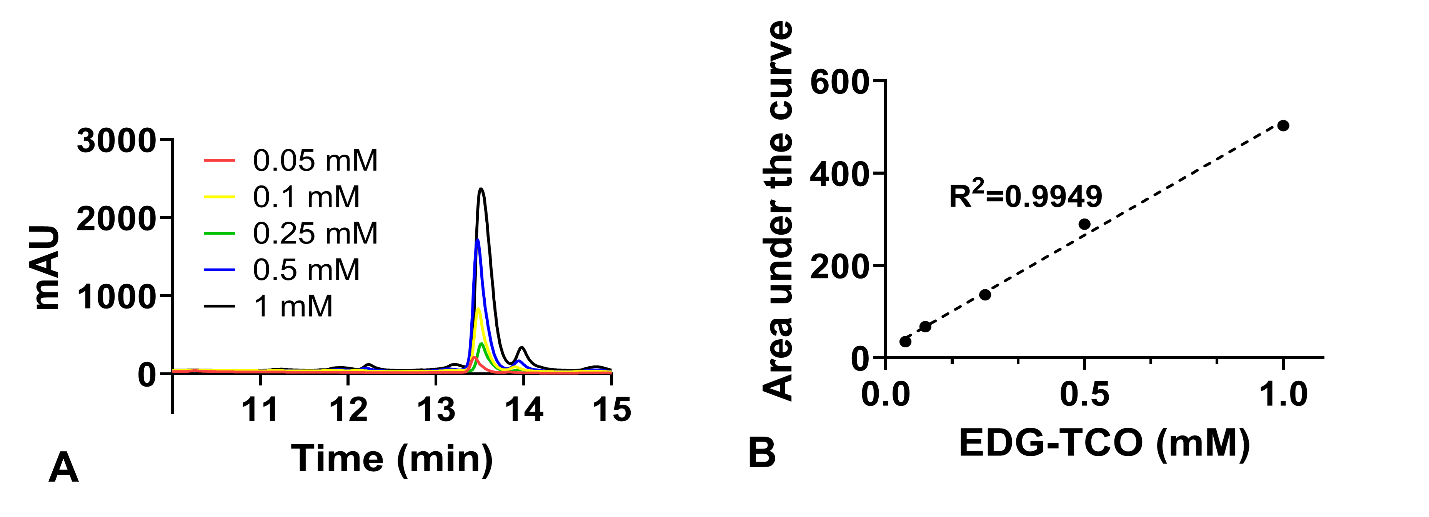

**Fig. S6:** Characterization of EDG retention in BOHAGel. (**A**) EDG-TCO was detected at 215 nm by HPLC. (**B**) The standard curve was developed using the peak area under the HPLC curve.

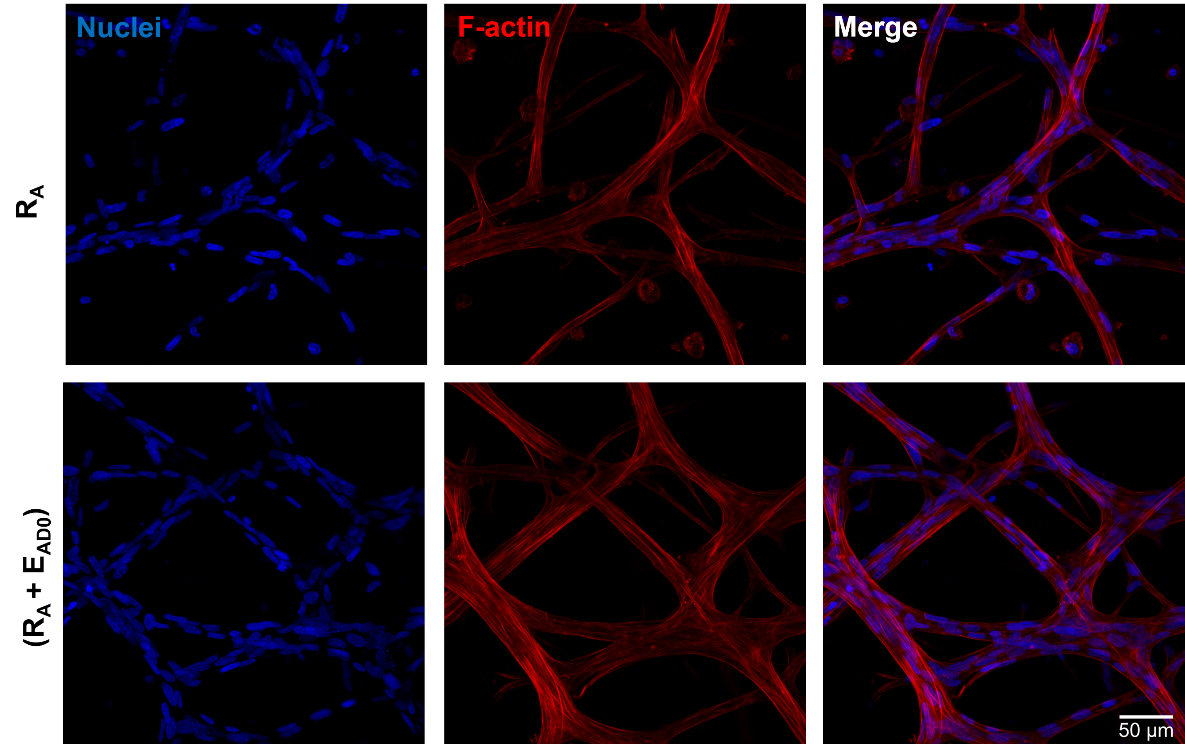

**Fig. S7:** Representative confocal images (25 ×) of NHLFs after 15 days of culture in BOHAGel under R_A_ and R_A_+E_AD0_ conditions. F-actin and cell nuclei were stained red and blue, respectively. Scale bar: 50 μm.

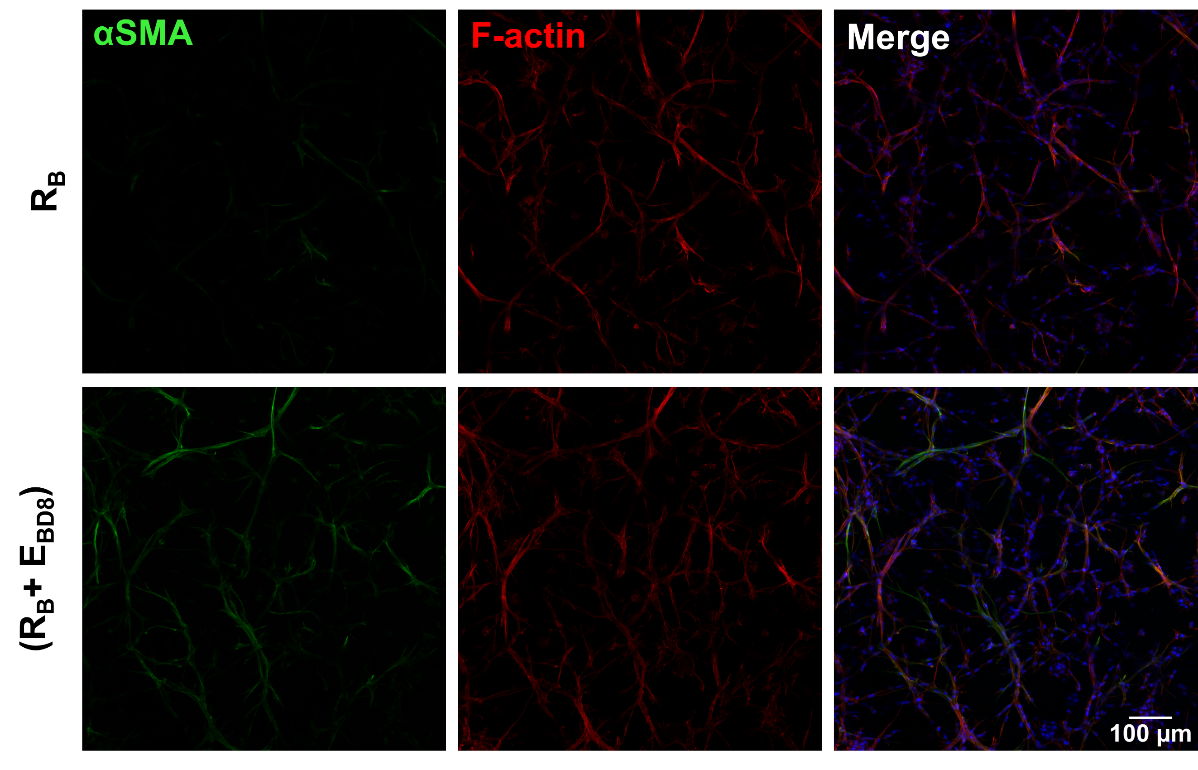

**Fig. S8:** Representative confocal images (10 ×) of NHLFs after 22 days of culture in BOHAGel under R_B_ and R_B_+E_BD8_ conditions. F-actin, αSMA, and cell nuclei were stained red, green, and blue, respectively. Scale bar: 100 μm.

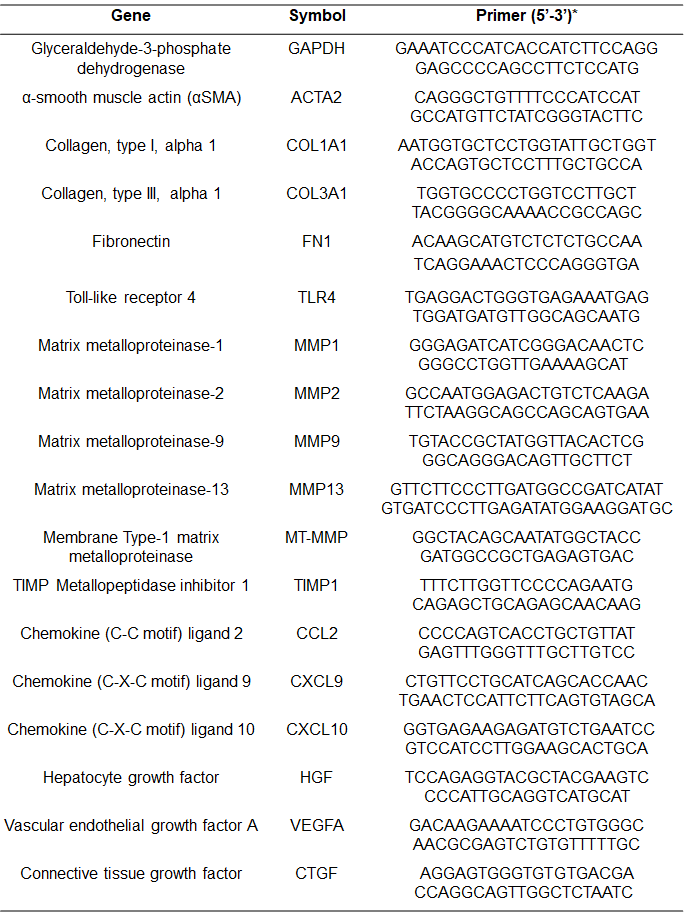

**Table S1:** List of primers used in qPCR analysis. *top: forward primer; bottom: reverse primer.

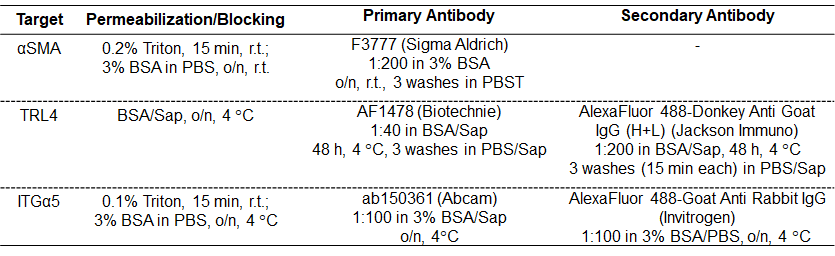

**Table S2:** Antibodies and conditions used for immunostaining. r.t.: room temperature; o/n: overnight. BSA/Sap: 0.05% (w/v) saponin in 3% (w/v) BSA in PBS**;** PBS/Sap: 0.05% (w/v) saponin in PBS.

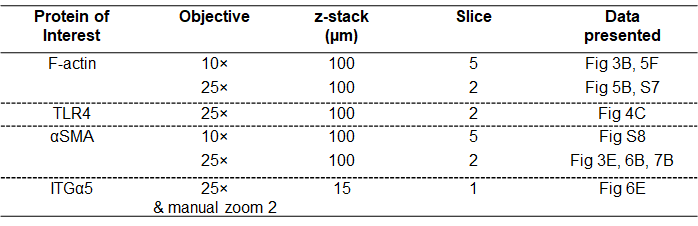
**Table S3:** Imaging acquisition settings for proteins of interest.

**Image Processing and Analysis:** Confocal z-stacks were converted to maximum intensity projections (MIP) and processed identically for each channel within each biological replicate. Background subtraction was performed using a rolling ball algorithm (radius = 50 pixels), followed by Gaussian blur (σ = 1.0 pixel) to reduce noise. A binary mask was generated from the F-actin channel using Otsu’s automated thresholding method, and this mask was used to define the cellular region of interest. For αSMA, the integrated fluorescence intensity (IFI) within this mask was measured and normalized to the integrated F-actin intensity within the same mask.^2,3^ For ITGα5, the IFI within the mask was divided by the mask area to obtain the mean fluorescence intensity (MFI). For the overlap coefficient, a binary αSMA mask was generated by applying Otsu’s automated thresholding method to the αSMA channel within the cellular region of interest. The αSMA mask was then applied to the F-actin channel, and the integrated F-actin intensity within the αSMA mask was divided by the total integrated F-actin intensity within the cellular region of interest. The resulting overlap coefficients correspond to the fraction of total F-actin intensity overlapping with αSMA-positive pixels.^2,3^
